# The C-terminal tail of polycystin-1 suppresses cystic disease in a mitochondrial enzyme-dependent fashion

**DOI:** 10.1101/2021.12.21.473680

**Authors:** Laura Onuchic, Valeria Padovano, Giorgia Schena, Vanathy Rajendran, Ke Dong, Nikolay P. Gresko, Xiaojian Shi, Hongying Shen, Stefan Somlo, Michael J. Caplan

## Abstract

Autosomal dominant polycystic kidney disease (ADPKD) is the most prevalent potentially lethal monogenic disorder. Approximately 78% of cases are caused by mutations in the *PKD1* gene, which encodes polycystin-1 (PC1). PC1 is a large 462-kDa protein that undergoes cleavage in its N and C-terminal domains. C-terminal cleavage produces fragments that translocate to mitochondria. We show that transgenic expression of a protein corresponding to the final 200 amino acid residues of PC1 in a *Pkd1*-KO orthologous murine model of ADPKD dramatically suppresses cystic phenotype and preserves renal function. This suppression depends upon an interaction between the C-terminal tail of PC1 and the mitochondrial enzyme Nicotinamide Nucleotide Transhydrogenase. This interaction modulates tubular/cyst cell proliferation, the metabolic profile, mitochondrial function and the redox state. Together, these results suggest that a short fragment of PC1 is sufficient to suppress cystic phenotype and open the door to the exploration of gene therapy strategies for ADPKD.

## INTRODUCTION

Autosomal dominant polycystic kidney disease (ADPKD) is the most common life-threatening monogenic disease, with a prevalence of ∼1:1000 (Lanktree et al., 2018). It is characterized by the progressive development of fluid-filled cysts whose expansion compromise renal function and can lead to end-stage renal disease. Mutations in the *PKD1* gene, which encodes polycystin-1 (PC1), are responsible for ∼78% of cases (Cornec-Le Gall et al., 2019). Although *PKD1* was identified over 25 years ago (Consortium, 1994), the downstream pathways leading to cystogenesis have not been fully elucidated, and current therapeutic options remain scarce and only modestly effective.

Over the past decade, metabolic abnormalities have emerged as a hallmark of ADPKD (Padovano et al., 2018). The first evidence for metabolic alterations in ADPKD was the observation of increased glycolysis and lactate production in cells from a *Pkd1* knockout (KO) mouse model (Rowe et al., 2013). Limiting glucose availability decreased their proliferation, and treatment with 2-deoxyglucose (2DG) to inhibit glycolysis led to partial amelioration of the cystic phenotype in *Pkd1*-KO mice (Rowe et al., 2013, Chiaravalli et al., 2016). The dependence on glycolysis and the increased lactate levels are together suggestive of an aerobic glycolysis phenotype, similar to the Warburg effect observed in cancer cells (Priolo and Henske, 2013, Rowe et al., 2013, Chiaravalli et al., 2016). Subsequently, other related significant metabolic alterations have been observed in ADPKD cellular and animal models, such as defective fatty acid oxidation and decreased rates of oxidative phosphorylation (Padovano et al., 2017, Hajarnis et al., 2017, Menezes et al., 2016). Several promising therapeutic approaches that target these altered metabolic pathways have been explored in animal models and produced phenotypic benefits, including anti-miR-17 or fenofibrate leading to increased levels of PPARα (Hajarnis et al., 2017), as well as metformin (Takiar et al., 2011), food restriction (Warner et al., 2016) and ketogenic diet (Torres et al., 2019). The mechanisms through which reduced or absent PC1 function produces metabolic alterations, however, have yet to be fully worked out. Recent data suggest the intriguing possibility that direct physical linkages between the PC1 protein and components of the mitochondrion may regulate mitochondrial bioenergetics (Padovano et al., 2017).

PC1 is an extremely large 462-kDa membrane glycoprotein that undergoes cleavage at its N- and C-termini (Padovano et al., 2020). The N-terminal cleavage occurs at the G protein-coupled receptor Proteolytic Site (GPS), giving rise to a 3048-amino acid (aa) N-terminal fragment (NTF) that remains non-covalently attached to the 1254-aa C-terminal fragment (CTF), which comprises PC1’s 11-transmembrane domains and its cytoplasmic tail (Qian et al., 2002). PC1 C-terminal cleavage generates several shorter PC1 CTF fragments and C-terminal tail fragments (PC1-CTT). An ∼100-kDa transmembrane CTF localizes to the endoplasmic reticulum where it may modulate store-operated calcium entry (Woodward et al., 2010). Several PC1-CTT fragments ranging from 17 to 34-kDa have been reported to translocate to the nucleus and to mitochondria (Chauvet et al., 2004, Lin et al., 2018, Low et al., 2006). Of note, expression of a protein construct corresponding to the 17-kDa PC1-CTT fragment rescued the fragmented mitochondrial network detected in cells lacking PC1 expression (Lin et al., 2018). This same study revealed that PC1-CTT, when expressed in *Drosophila melanogaster*, leads to increased CO_2_ production and decreased capacity for endurance exercise (Lin et al., 2018). While these findings suggest that the PC1-CTT might alter mitochondrial function *in vivo*, the effects of re-expressing PC1-CTT in a PC1-deficient vertebrate model of ADPKD have not been assessed.

We expressed the PC1-CTT in a *Pkd1*-KO ADPKD mouse model and characterized the resultant phenotype. We show that PC1-CTT expression significantly reduces the severity of cystic disease and that this effect is dependent upon interaction between the PC1-CTT and the mitochondrial enzyme Nicotinamide Nucleotide Transhydrogenase (NNT). Expression of PC1-CTT in NNT-deficient mice does not produce disease amelioration. Furthermore, we show that PC1-CTT re-expression in the presence of NNT leads to increased mitochondrial mass, altered redox modulation, increased assembly of ATP synthase at a “per mitochondrion” level, and decreased tubular epithelial cell proliferation, suggesting potential mechanisms for the observed rescue. Similarly, unbiased metabolomics reveals that the ability of the PC1-CTT to normalize the *Pkd1*-deficient metabolic profile is consistent with PC1-CTT modifying the normal function of NNT. Finally, we show that PC1-CTT re-expression in *Pkd1-*KO mice can rescue NNT enzymatic activity, as measured *ex vivo*, to the level observed in healthy wild type (WT) controls. Our discovery that a short fragment of the PC1 protein is sufficient to substantially reduce disease severity in a mouse model opens the door to exploration of gene therapy strategies for the treatment of ADPKD.

## RESULTS

### Expression of PC1-CTT suppresses cystic phenotype in an orthologous murine model of ADPKD

We generated a BAC construct in which the sequence of the PC1-CTT (aa 4102-4302; the final 200 residues of human PC1) with a 2XHA epitope tag added at its N terminus (CTT)(Merrick et al., 2019) is inserted into the Rosa26 locus. This construct includes a neomycin resistance (NeoR) STOP cassette flanked by loxP sequences upstream of the CTT sequence so as to permit the CTT protein to be produced only in cells that express Cre. A transgenic mouse line with the BAC transgene stably incorporated into its germline was generated and crossed with a previously characterized orthologous, conditional *Pkd1*-KO mouse model of ADPKD (*Pkd1*^fl/fl^; Pax8^rtTA^; TetO-Cre) (Ma et al., 2013). Doxycycline induction from p28-p42 of these second-generation mice (2HA-PC1-CTT; *Pkd1*^fl/fl^; Pax8^rtTA^; TetO-Cre) on the C57BL/6N (“N”) background leads to CTT expression in renal epithelial cells that have undergone Cre-driven disruption of *Pkd1* and therefore do not produce functional PC1 protein (Figure 1A). We generated cohorts of littermates with comparable sex distributions that did or did not carry the CTT BAC transgene on the “N” background (N-*Pkd1*-KO+CTT vs N-*Pkd1*-KO mice) and evaluated them 10 weeks after completion of induction at 16 weeks of age (Figures 1B-1E). The mice that expressed the CTT presented significant reduction of their cystic burden. These N-*Pkd1*-KO+CTT animals had a 60% lower kidney-to-body weight (KW/BW) ratio than did the N-*Pkd1*-KO animals that did not inherit the transgene (Figure 1C). Consistent with the preservation of renal size and gross morphology observed in the CTT expressing animals, these mice exhibited a significant reduction in blood urea nitrogen (BUN) and serum creatinine, revealed by a 3 and 3.5-fold change, respectively, in comparison to the N-*Pkd1*-KO littermates that did not inherit the transgene. Of note, the magnitudes of these 3 parameters did not significantly differ between N-*Pkd1*-KO+CTT animals and healthy WT controls.

**Figure 1:**
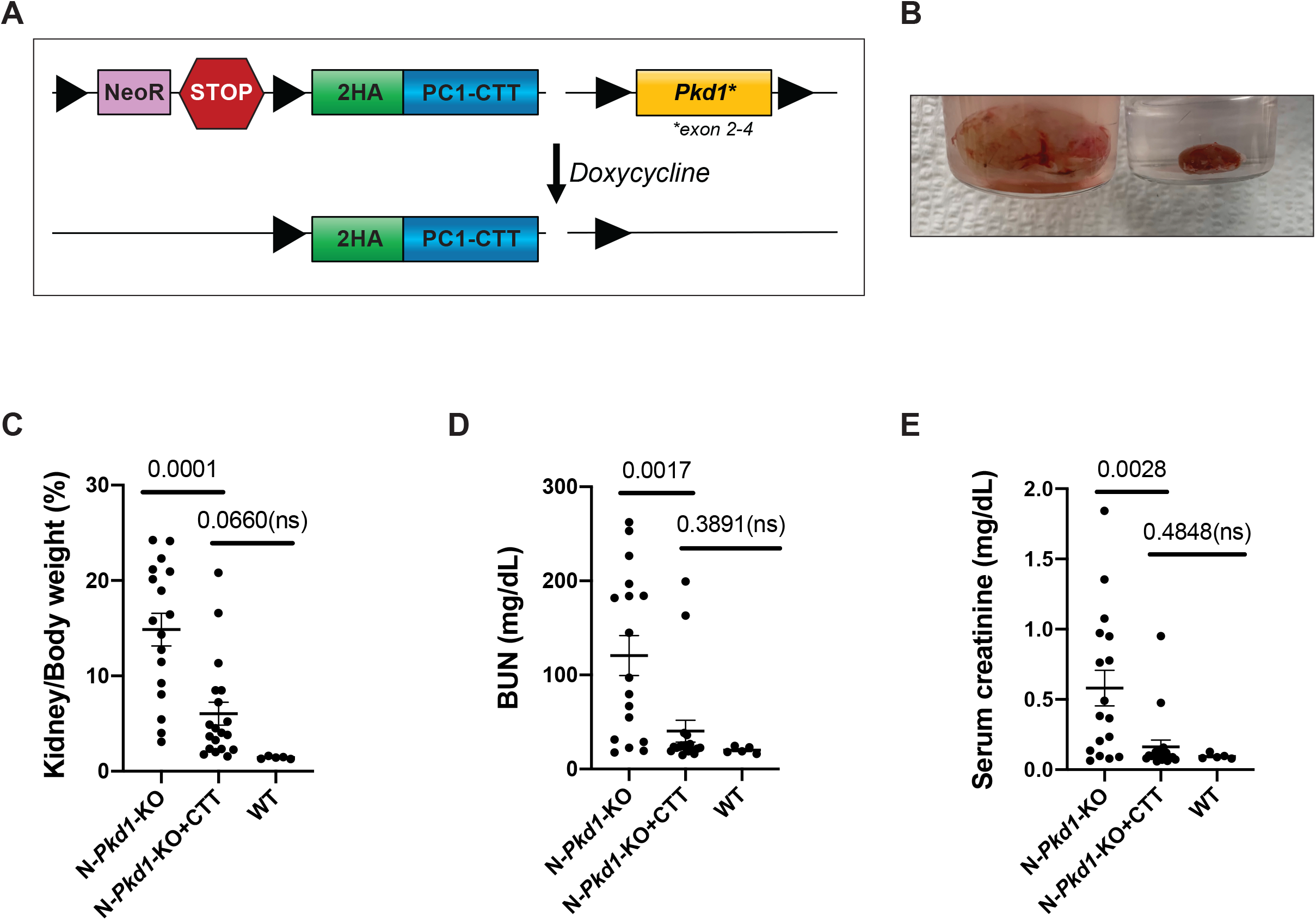
Expression of polycystin-1 C-terminal tail (CTT) suppresses cystic disease in an orthologous mouse model of ADPKD. (A) Design of the 2HA-PC1-CTT; *Pkd1*^fl/fl^; Pax8^rtTA^; TetO-Cre (*Pkd1*-KO+CTT) mouse model. Briefly, exons 2-4 of the *Pkd1* gene are flanked by loxP sequences. These mice also carry a BAC-2HA-PC1-CTT transgene inserted in the Rosa26 locus with a *loxP-*STOP-*loxP* cassette upstream of it. Cre-mediated recombination via doxycycline induction promotes 2HA-PC1-CTT expression and loss of full-length PC1 in tubular epithelial cells. (B) Gross mouse kidney anatomy from two induced female littermates, in which one animal did not inherit the BAC CTT transgene (left), and the other inherited the transgene and exhibits a profound suppression of the cystic phenotype (right). (C-E) Comparative analysis between N-*Pkd1*-KO+CTT and N-*Pkd1*-KO mice showing differences in KW/BW ratio (C), BUN (D) and serum creatinine (E). Cystic mouse cohorts are composed of 53%-58% female and 42%-47% male mice. Data are expressed as mean ± SEM. Pairwise comparisons were performed using Student’s t-test. See also Figure S1.

We next wished to assess whether the level of CTT expression in the BAC transgenic mice is comparable to the level of PC1-CTT that is produced through cleavage of the full length PC1 protein in mouse renal epithelial cells *in vivo*. Full-length WT PC1 is a low-abundance protein that is not reliably detectable in mouse kidneys (Lin et al., 2018, Cai et al., 2014). Not surprisingly, therefore, the endogenous cleavage products of PC1, including the PC1-CTT, are even less abundant and also not detectable by standard methods. To address this problem, we employed the previously characterized full-length *Pkd1*^F/H^-BAC transgenic mouse line (BAC-*Pkd1*) (Fedeles et al., 2011, Cai et al., 2014) that expresses a tagged version of the PC1 protein under the control of its native promoter. The BAC-encoded PC1 protein is tagged at its N-terminus with a triple-FLAG epitope inserted after the signal sequence and at its C-terminus with a triple-HA sequence inserted prior to the stop codon (3FLAG-PC1-3HA) (Figure 2A) (Fedeles et al., 2011). Our study made use of offspring of a founder of the *Pkd1*^F/H^-BAC line (Tg248) that carries 3 copies of the BAC-*Pkd1* transgene. Quantitative western blotting has previously determined that Tg248 mice exhibit a 3-fold increase in tagged PC1 expression relative to offspring of a single-copy founder (Tg14) (Cai et al., 2014). Since expression of tagged PC1 in the Tg248 line is driven by the complete, endogenous *Pkd1* promoter, it is likely that the 3 copies of the *Pkd1*^F/H^-BAC transgene together drive expression of the tagged protein that is roughly comparable to 1.5X the quantity of the native PC1 generated from the 2 native copies of the *Pkd1* gene encoded in the mouse genome. We performed quantitative western blotting on 60 μg of lysates prepared from kidneys of doxycycline induced N-*Pkd1*-KO+CTT mice and from kidneys of *Pkd1*^F/H^-BAC mice. This analysis revealed similar CTT protein expression levels in N-*Pkd1*-KO+CTT mice when compared to the expression of the equal molecular weight PC1-CTT-3HA that results from the cleavage of full-length 3FLAG-PC1-3HA in *Pkd1*^F/H^-BAC mice (Figure S1). This observation suggests an upper threshold for CTT expression in our N-*Pkd1*-KO+CTT mice of approximately 1.5-fold above the levels expected for WT animals. Thus, the rescue of the cystic phenotype that is observed in the N-*Pkd1*-KO+CTT mice is not a consequence of massive overexpression of super-physiological levels of the CTT.

**Figure 2:**
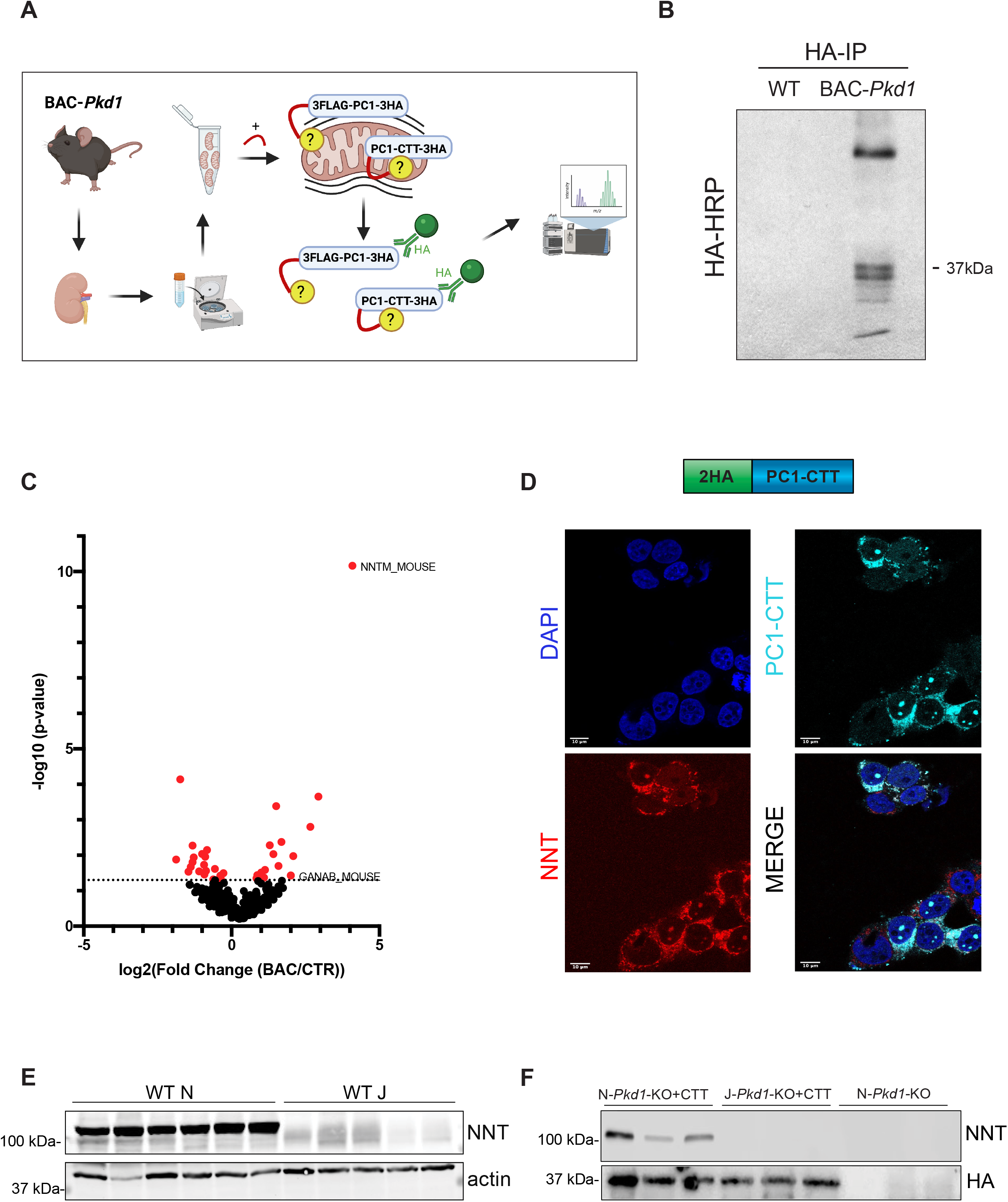
2HA-PC1-CTT (CTT) colocalizes and interacts with the mitochondrial enzyme NNT. (A) Schematic representation of the workflow for the identification of protein interaction partners from mitochondrial-associated pools of PC1 and its fragments. Crude mitochondria fractions prepared from *Pkd1*^F/H^-BAC and WT mice were solubilized, crosslinked with 3mM DTSSP, and subjected to immunoprecipitation with anti-HA antibodies. The proteins recovered were identified by mass spectrometry. (B) HA-HRP immunoblotting of the crude mitochondrial HA-immunoprecipitate revealed recovery of both full-length 3FLAG-PC1-3HA and PC1-CTT-3HA fragments only in immunoprecipitates from *Pkd1*^F/H^-BAC mice. (C) Volcano plot of PC1 and PC1-CTT interacting partners from *Pkd1*^F/H^-BAC vs WT kidneys (*P*<0.05 for colored dots determined by Fisher’s exact test; n=3 per group). (D) Representative immunofluorescence (63X) image showing transient transfection of the 2HA-PC1-CTT construct in HEK293 cells, revealing mitochondrial colocalization between the construct, identified with anti-PC1-C-terminus antibodies, and endogenous NNT. (E) Immunoblotting of total kidney lysate from WT “N” and “J” mice, confirming the presence and absence of NNT, respectively. (F) Immunoblotting of HA immunoprecipitate, revealing immunoprecipitation of CTT in both “N” and “J” *Pkd1*-KO+CTT mice. NNT coimmunoprecipitation is detected exclusively in immunoprecipitate from N-*Pkd1*-KO+CTT mice. Scale bar:10 μm. See also table S1.

### PC1-CTT interacts and colocalizes with the mitochondrial enzyme Nicotinamide Nucleotide Transhydrogenase (NNT)

We undertook studies designed to identify the downstream pathways linking PC1-CTT re-expression and disease amelioration. To do so, we again employed the *Pkd1*^F/H^-BAC mouse model, which is generated on a mixed strain background (Fedeles et al., 2011). Previous data showed that full-length PC1 localizes to mitochondria-associated endoplasmic reticulum membranes (MAMs)(Padovano et al., 2017), while PC1-CTT can localize to mitochondrial matrix (Lin et al., 2018). We sought to identify possible protein interaction partners of these mitochondria-associated pools of PC1 and its fragments. *Pkd1*^F/H^-BAC mouse kidneys were homogenized and a crude mitochondria fraction was isolated by differential centrifugation (Wieckowski et al., 2009). This fraction was expected to contain MAM-associated full-length PC1 and PC1-CTT cleavage products within the recovered mitochondria. Following addition of the covalent crosslinking reagent DTSSP, mitochondria were solubilized and the lysate was subjected to immunoprecipitation using magnetic beads coupled to anti-HA antibodies (Figure 2A). Immunoblot analysis of this precipitate employing an HRP-conjugated anti-HA antibody revealed not only the presence of the 150-kDa CTF derived from full-length 3FLAG-PC1-3HA protein, but also 17-37-kDa PC1-CTT-3HA fragments (Figure 2B). None of these polypeptides were detected in immunoprecipitates from WT control animals. Next, in order to identify putative PC1 and PC1-CTT interactors, we performed mass spectrometric proteomic analysis of the material immunoprecipitated from *Pkd1*^F/H^-BAC mice and compared the results to a mass spectrometric analysis performed on immunoprecipitates from WT controls. Potential PC1 interactors identified in this way were cross-referenced against the MitoCarta 2.0 mitochondrial proteome (Calvo et al., 2016). The most significantly enriched interactor identified was the mitochondrial enzyme Nicotinamide Nucleotide Transhydrogenase (NNT) (Figure 2C and Table S1). The amount of NNT detected in *Pkd1*^F/H^-BAC kidney immunoprecipitates was >16-fold greater than that detected in control WT immunoprecipitates (*P*≤10^-10^). Of note, we identified other interesting PC1-relevant hits specific to the endoplasmic reticulum (ER) proteome, consistent with the fact that the crude mitochondria fractions contain MAMs. This is the case for the observed 8-fold enrichment of *Ganab* (*P*≤0.05), which encodes for glucosidase II subunit α and that, when mutated, can lead to ADPKD (Porath et al., 2016, Besse et al., 2017).

We next sought to assess the validity of the putative interaction between PC1 and NNT. Because the catalytic domain of NNT localizes to the matrix-facing leaflet of the mitochondrial inner membrane, we expected that any biologically-relevant interaction between NNT and PC1 involved the PC1-CTT rather than the intact, full-length PC1. Due to the low expression level of CTT in our mouse models, it was not possible to reliably assess its subcellular localization through immunofluorescence microscopy. To assess whether the CTT distribution was consistent with its interaction with NNT, we transiently transfected WT HEK293 cells with the 2HA-PC1-CTT construct and found that it colocalized with endogenous NNT at the mitochondria (Figure 2D). Of note, CTT expression was also observed in nuclei of transfected cells, consistent with previous findings (Chauvet et al., 2004).

Interestingly, the widely used C57BL/6J (“J”) mouse strain carries a deletion of exons 7-11 in the *Nnt* gene, a mutation that completely abrogates its expression (Figure 2E). This variant was acquired prior to 1984, however was only identified in 2005, when it was associated with glucose intolerance in “J” mice (Toye et al., 2005). We moved the alleles required to produce the *Pkd1*-KO+CTT mice to this NNT-deficient background (J-*Pkd1*-KO+CTT). To determine whether NNT interacts with 2HA-PC1-CTT *in vivo*, we performed anti-HA pulldowns from N-*Pkd1*-KO+CTT, J*-Pkd1*-KO+CTT and N-*Pkd1*-KO total kidney lysates. We found that NNT is only detected in immunoprecipitates derived from animals that express the 2HA-tagged PC1-CTT and not in those derived from N-*Pkd1*-KO mice that do not express CTT (Figure 2F). As expected, anti-HA immunoprecipitates from *Pkd1*-KO+CTT kidneys on the “J” background did not contain a 114-kDa NNT band that could be detected on immunoblotting with anti-NNT antibody. Taken together, these data demonstrate that the PC1-CTT interacts with the inner mitochondrial membrane protein NNT in mouse kidney epithelial cells *in vivo*.

### The PC1-CTT/NNT interaction modulates disease progression

We next employed the J-*Pkd1*-KO+CTT mice to evaluate the relevance of the PC1-CTT/NNT interaction to disease progression. While the expression of CTT on the “N” background leads to significant phenotype amelioration, no reduction in KW/BW ratio was observed on the “J” background when comparing J-*Pkd1*-KO+CTT mice to J-*Pkd1*-KO littermates that did not inherit the CTT-encoding BAC transgene (Figure 3A). Similarly, BUN and serum creatinine levels were not significantly different between the two groups (Figures 3B and 3C). To further characterize these models, we measured tubular and cystic area relative to whole kidney area. This parameter is significantly smaller in *Pkd1*-KO+CTT vs *Pkd1*-KO mice on the “N” background (Figure 3D). In marked contrast, no difference was observed when performing the same comparison with mice that manifest these genotypes on the “J” background. Additionally, we performed immunofluorescence microscopy on renal tissue from mice of all four models and quantitated the extent of tubular epithelial cell proliferation by determining the fraction of Ki67-positive nuclei. While the fraction of Ki67-positive nuclei was reduced by a factor of 2.3 in *Pkd1*-KO+CTT compared to *Pkd1*-KO mice on the “N” background, proliferation levels remained equivalent and elevated in these same models on the “J” background (Figures 3E and 3F). Of note, genotyping for the presence of WT *Nnt* (“N” mice) and mutant *Nnt* (“J” mice) was performed at p6-p12 and confirmed upon sacrifice. Phenotype amelioration by CTT expression, therefore, requires the presence of NNT (Figure 3G), suggesting that the ability of the CTT to suppress cystic disease is dependent upon the PC1-CTT/NNT interaction.

**Figure 3:**
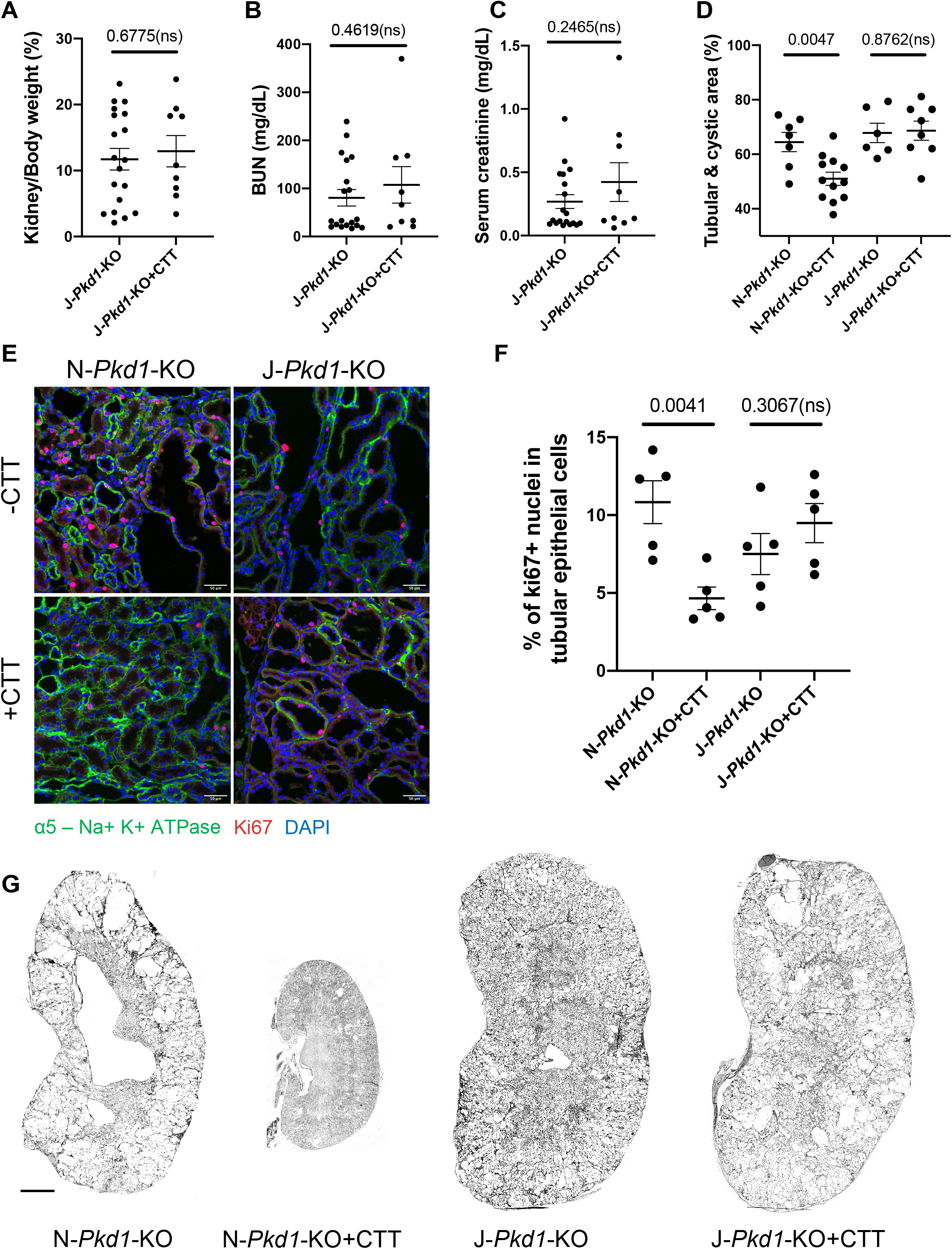
The interaction between NNT and CTT is critical in preventing cystogenesis and tubular proliferation. (A-C) Comparative analysis between *Pkd1*-KO+CTT and *Pkd1*-KO in the “J” background revealed no significant change in KW/BW ratio (A), BUN (B) and serum creatinine levels (C). Cystic mouse cohorts are composed of 53%-55% female and 45%-47% male mice. (D) Quantification of tubular and cystic area in H&E-stained kidney sections from *Pkd1*-KO and *Pkd1*-KO+CTT mice on both “N” and “J” backgrounds, as determined by ImageJ using images of renal cross-sections depicted in (G). (E) Representative immunofluorescence images (20X) showing tubular proliferation in *Pkd1*-KO and *Pkd1*-KO+CTT mice on “N” and “J” backgrounds, assessed by ki67 staining. Tubular epithelial cells are identified by positive Na,K-ATPase α-subunit staining. Scale bar: 50 μm (F) Quantification of tubular proliferation determined by the percentage of ki67 positive nuclei (n= 9 images per mouse, 5 mice per group). Counting of ki67 positive nuclei was performed by an individual blinded to the experimental conditions. (G) H&E-stained kidney sections (4X) from 16-week *Pkd1*-KO and *Pkd1*-KO+CTT mice on the “N” and “J” backgrounds. Scale bar: 200 μm Data are expressed as mean ± SEM. Pairwise comparisons were performed using Student’s t-test. See also Figures S2, S3 and S4.

To ensure that no other factors might contribute to the different responses of the “N” and “J” strains to CTT expression, we assessed whether there were any systematic differences in the level of Cre-expression between these mouse models. In the *Pkd1^f^*^l/fl^; Pax8^rtTA^; TetO-Cre model, oral doxycycline induces activation of TetO-Cre under the control of the Pax8^rtTA^ promoter, leading to excision of the floxed exon 2-4 region and consequent inactivation of *Pkd1* in Pax8^rtTA^-exppressing cells (Shibazaki et al., 2008, Ma et al., 2013). We determined the status of homozygosity or heterozygosity for both the Pax8^rtTA^ and TetO-Cre alleles by quantitative PCR (qPCR) for each animal included in the present cohort and found that Pax8^rtTA^ and TetO-Cre copy numbers were randomly distributed across the 4 groups (*Pkd1*-KO+CTT and *Pkd1*-KO in both “N” and “J” backgrounds), and did not correlate to any degree with disease severity, as determined by KW/BW ratio (Figure S2). We also evaluated the efficiency of *Pkd1* knock-out in all animals to determine whether there was any significant genotype-dependent effect on this parameter. We determined levels of non-rearranged WT *Pkd1* by extracting genomic DNA from kidney tissue from each mouse contained in the cohort followed by quantitative genomic PCR using a reverse primer specific for *Pkd1* exon 4 and a forward primer specific to its preceding intron. We calculated a relative ratio of non-rearranged *Pkd1* by comparing the observed levels of non-rearranged *Pkd1* product to the levels obtained from WT control kidneys and found that rearrangement levels are equivalent across all 4 mouse groups (Figure S3). Finally, the NNT mutation appears to be the major allelic difference between C57BL/6J and C57BL/6N strains (Simon et al., 2013) and is the only candidate genetic variation that has been directly associated with the metabolic (Toye et al., 2005, Ronchi et al., 2013, Fergusson et al., 2014), cardiologic (Murphy, 2015) and renal (Usami et al., 2018) differences observed between them. It is important to note, however, that the “N” and the “J” strains manifest at least one additional phenotypically significant genetic polymorphism. This is the case for the *Crb1* gene, in which the *rd8* retinal degeneration mutant is detected exclusively in the “N” mice and results in a recessive ocular phenotype. To our knowledge, this is the only other mutation that differs between “N” and “J” mice and that is directly associated to a clear phenotype (Mattapallil et al., 2012). We expected that the *rd8* mutation would be present in both N-*Pkd1*-KO+CTT and N-*Pkd1*-KO mice. We genotyped the “N” animals in the cohort for this mutation and show comparable *rd8* allele distribution among both groups (Figure S4), thus excluding the possibility that skewed distributions of this mutant allele that were inadvertently introduced during the breeding of our mouse colonies could account for the observed phenotypic difference between both groups. Furthermore, this observation suggests that any other potential variants should have a similar distribution across both groups and are unlikely to be responsible for the observed phenotype differences.

### PC1-CTT expression produces a change in metabolic profile in the presence of NNT

In light of the localization of the 2HA-PC1-CTT to mitochondria and its NNT-dependent effects on the progression of cystic disease, we sought to evaluate potential metabolic consequences of CTT expression in our *Pkd1*-KO mice. We performed Liquid-Chromatography Mass-Spectrometry (LC-MS) based metabolite profiling on whole kidney tissue extracts across all 4 experimental groups 10 weeks after the end of the doxycycline induction of Cre expression. While Principal Component Analysis (PCA) and hierarchical clustering revealed a distinct separation between *Pkd1*-KO and *Pkd1*-KO+CTT mice on the “N” background, it failed to distinguish among these two groups on the “J” background (Figure 4A). Unpaired *t-test* resulted in the detection of 47 metabolites that significantly changed (*P* <0.05) between *Pkd1*-KO+CTT and *Pkd1*-KO mice on the “N” background. There were 44 identified metabolites that met the criteria of both *P* value <0.05 and fold change >2, of which 6 were upregulated and 38 were downregulated in N-*Pkd1*-KO+CTT compared to N-*Pkd1*-KO kidneys (Figure 4B and Table S2). In contrast, analysis of both J-*Pkd1*-KO+CTT and J-*Pkd1*-KO only revealed a significant change (*P* value <0.05 and fold change >2) in 1 metabolite.

**Figure 4:**
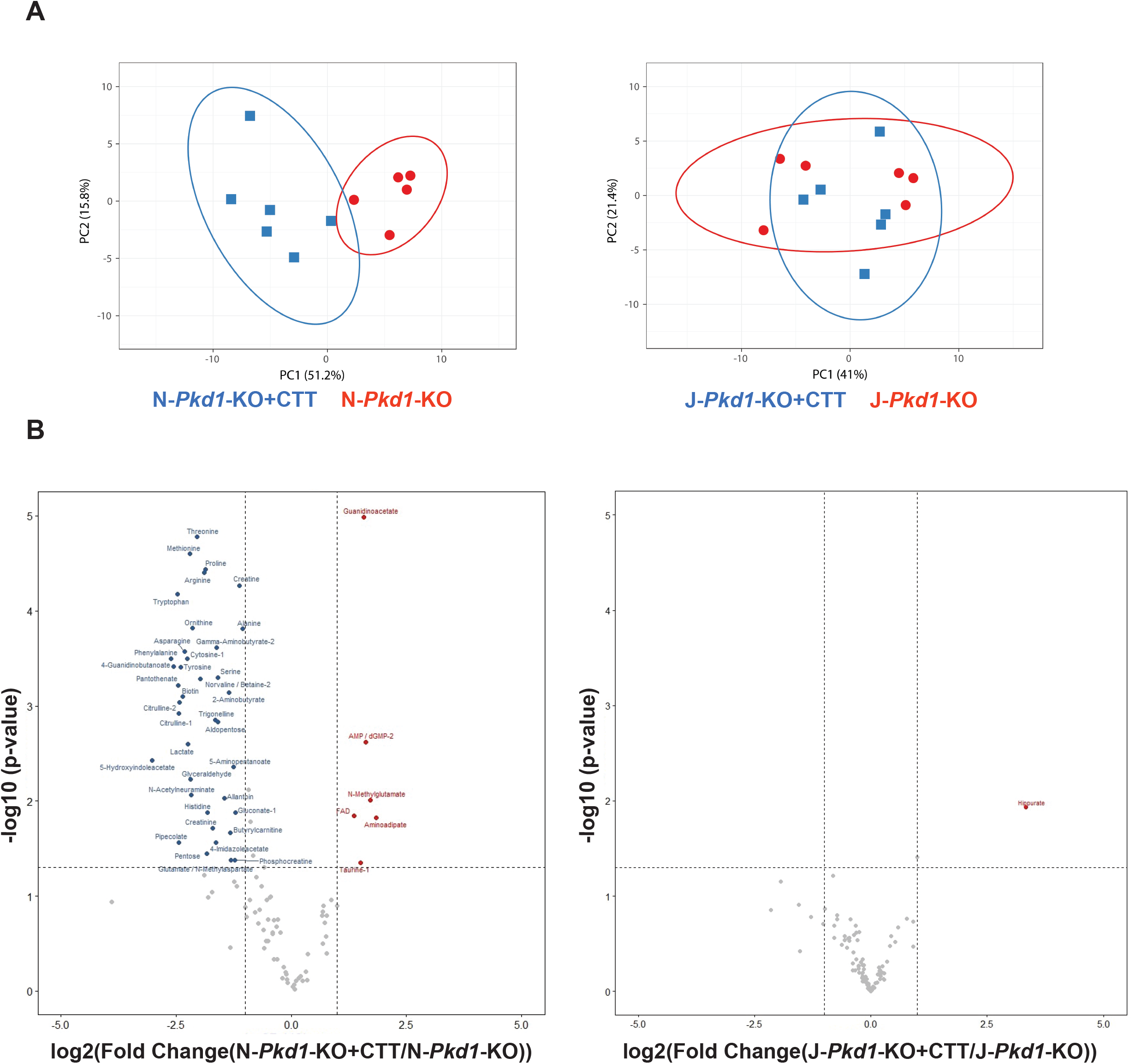
CTT expression produces a change in metabolic profile in the presence of NNT. (A) PCA plot of metabolomic data from *Pkd1*-KO+CTT mice vs *Pkd1*-KO mice revealed a clear separation between groups in the “N” background but not in the “J” background. Each individual mark corresponds to a different sample (n=5 or 6 mice per group) and its location in the plot is determined by a linear combination of metabolites. (B) Volcano plot showing metabolite profiling of the kidney extracts from *Pkd1*-KO+CTT mice vs *Pkd1*-KO mice in the “N” and “J” backgrounds. The vertical lines in each panel mark 2-fold changes. The horizontal lines mark *P*<0.05; determined by Student’s t-test. Labels and colored dots indicate metabolites with significant fold changes. See also Table S2.

Interestingly, many of the metabolites whose levels are reduced in N-*Pkd1*-KO+CTT mice have been previously implicated in ADPKD pathogenesis and some of them are related to potential therapeutic targets (Figure 4B). This is the case for the 4-fold decrease in methionine levels induced by CTT expression in the “N” mice. Methionine has been reported to promote cyst growth through increased Mettl3 expression and to constitute a potential therapeutic vulnerability in murine models (Ramalingam et al., 2021). Similarly, we report that CTT expression in N mice is associated with a 4-fold decrease in lactate levels, consistent with a reduction in the dependence on glycolysis that has been detected in ADPKD cell and mouse models (Rowe et al., 2013, Podrini et al., 2018, Chiaravalli et al., 2016). Additionally, a recent study has found that *Pkd1*^-/-^ mouse embryonic fibroblasts are dependent upon glutamine to fuel the TCA cycle and that these cells exhibit increased asparagine synthetase activity (Podrini et al., 2018). This enzyme couples the conversion of aspartate-to-asparagine to the conversion of glutamine-to-glutamate. These findings are consistent with the results of plasma metabolomics in children and young adults with ADPKD relative to healthy controls, which revealed a significant increase in asparagine levels (Baliga et al., 2021). Interestingly, we detected a 5-fold decrease in asparagine levels and a 2.5-fold decrease in glutamate levels in the N-*Pkd1*-KO+CTT mice as compared to those detected in N-*Pkd1*-KO mice, suggesting a rescue of this feature of pathologic metabolic reprogramming. Finally, we observed significant reductions in the levels of known uremic toxins such as allantoin and 5-hydroxyindoleacetate in N-*Pkd1*-KO+CTT mice, as well as a significant reduction in the levels of the urea cycle metabolites ornithine, citrulline and arginine. Of note, the previously mentioned clinical metabolomic study (Baliga et al., 2021) suggests drastic changes in urea cycle activity in ADPKD and reveals a significant positive association between ornithine levels and height-adjusted total kidney volume (HtTKV) at baseline, as well as association between ornithine levels and change in HtTKV over a 3-year observation period. In conclusion, the metabolic signature associated with CTT expression in N-*Pkd1*-KO mice is marked by the reversal of dysregulated metabolites that are associated with ADPKD.

### N-*Pkd1*-KO+CTT mice exhibit increased NNT expression and increased assembly of ATP synthase and mitochondrial complex IV at a “per mitochondrion” level

We next sought to determine whether CTT and CTT/NNT interactions alter the inventories of proteins that potentially affect mitochondrial function. To this end, we performed immunoblotting analyses on kidney lysates from N-*Pkd1*-KO+CTT and N-*Pkd1*-KO mice (Figure 5A). We detected a 4-fold increase in NNT expression (normalized to actin) in CTT-expressing mice compared to N-*Pkd1*-KO littermates that did not inherit the transgene. Interestingly, this significant difference was shown to be a product of both increased mitochondrial mass in CTT expressing mice (as revealed by the significant 1.8-fold increase in TOMM20/actin ratio), as well as of increased NNT expression at a “per mitochondrion” level (as revealed by the significant 2.1-fold increase in NNT/TOMM20 ratio in these same mice) (Figure 5B). Comparable findings were identified in tissue from cystic human kidneys, which exhibit a significant decrease in NNT protein expression as compared to healthy kidney tissue (Figure 5D). Furthermore, western blotting employing a “mitococktail” antibody that interrogates the levels of stably assembled mitochondrial membrane complexes demonstrated increased levels of assembled ATP-synthase (complex V or CV) and cytochrome c oxidase (complex IV or CIV) at a “per mitochondrion” level, as revealed by a significant 2.4-fold increase in CV/TOMM20 and a 1.9-fold increase in CIV/TOMM20 in N-*Pkd1*-KO+CTT expressing mice (Figures 5A and 5B). Interestingly, we did not observe any differences in the assembly of complexes I (CI), II (CII) or III (CIII) in these same mice. A similar evaluation on the “J” background reveals no significant difference in mitochondrial mass (TOMM20/actin) or in mitochondrial complex assembly (Figures 5A and 5C), suggesting that the effects observed in the “N” background are specific to the CTT expression and the potential for its interaction with NNT.

**Figure 5:**
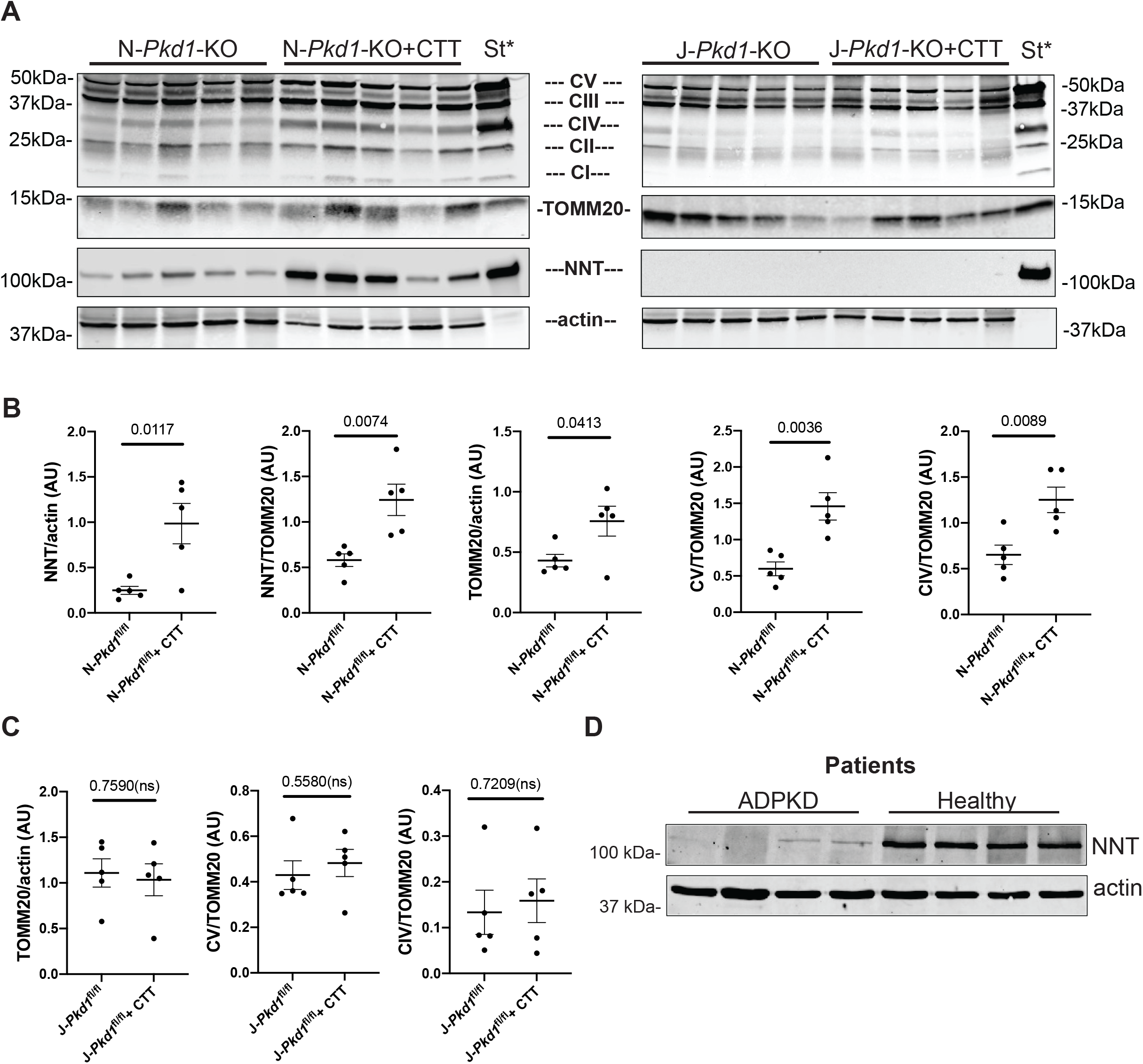
CTT expression in cystic “N” mice leads to increased NNT protein levels, mitochondrial mass and stable assembly of ETC complexes. (A-C) Immunoblots of total kidney lysate from *Pkd1*-KO+CTT and *Pkd1*-KO mice on the “N” and “J” backgrounds using a “mitococktail” antibody that reports on the assembly status of mitochondrial complexes I, II, III, IV and V. Blots were also probed with antibodies directed against TOMM20 and NNT. Blotting for actin served as a loading control (A). *St: mitochondrial extract from rat heart tissue lysate (catalog #ab110341, Abcam), serving as standard positive experimental control, indicates the positions of mitochondrial complexes I, II, III, IV and V. Individual bands were quantified and normalized to protein content (actin) or mitochondrial content (TOMM20). Normalized band intensities within the same membrane were compared and shown in graphs depicting the significant differences between N-*Pkd1*-KO+CTT vs N-*Pkd1*-KO mice (B) and the corresponding analyses in the “J” background (C). (D) Immunoblot depicting NNT protein levels in renal tissue from ADPKD and non-cystic human patients. Data are expressed as mean ± SEM. Pairwise comparisons were performed using Student’s t-test.

### NNT expression alone is not sufficient to suppress the cystic phenotype in ***Pkd1*-KO mice**

We next interrogated whether NNT alone, in the absence of CTT, serves as a significant gene modifier in the context of ADPKD. To that end, we compared data previously presented (Figures 1C-E and 3 A-C) derived from N-*Pkd1*-KO mice and J-*Pkd1*-KO to each other, in the absence of CTT expression. Surprisingly, NNT-competent cystic mice exhibited non-significant trends towards increased KW/BW ratio (Figure 6A) and BUN levels (Figure 6B), as well as a significant 2.16-fold increase in serum creatinine levels (Figure 6C) when compared to J-*Pkd1*-KO. Considering both mouse cohorts were analyzed at 16-weeks, a time window characterized by marked and advanced cystogenesis (Ma et al., 2013), it is not surprising that only the least sensitive parameter would show a statistically significant difference at this late stage of disease progression. It is well established that declining glomerular filtration rate (GFR) is a late consequence of cyst progression in ADPKD (Grantham et al., 2011) consistent with the observed difference in serum creatinine levels but not in other parameters that have already reached their maximal cystic disease-induced elevations. Furthermore, we performed immunohistochemistry (IHC) on mouse kidney tissue obtained from these *Pkd1*-KO mice on both backgrounds and from N-*Pkd1*-KO+CTT mice (Figure 6D). As expected, the “J” cystic mice did not express NNT. The cystic “N” mice, both in the presence and absence of CTT, expressed NNT predominantly in distal segments of the nephron and to a lesser degree in proximal tubules. No NNT was detected in glomeruli or Bowman’s capsule. A similar pattern of NNT renal distribution is reported in the Human Protein Atlas (proteinatlas.org)(Uhlen et al., 2015). It is important to note that CTT-expressing mice exhibit dramatic preservation of renal parenchyma architecture as compared to “J” and “N” mice lacking CTT expression. Finally, cystic mice on the “N” background were frequently found to exhibit mild to moderate hydronephrosis, characterized by an expanded pelvis (Figure 6E) with no obvious downstream obstruction and negative staining for calcium oxalate deposition (data not shown). Thus, a cause for this specific morphologic feature in “N” mice remains to be determined.

**Figure 6:**
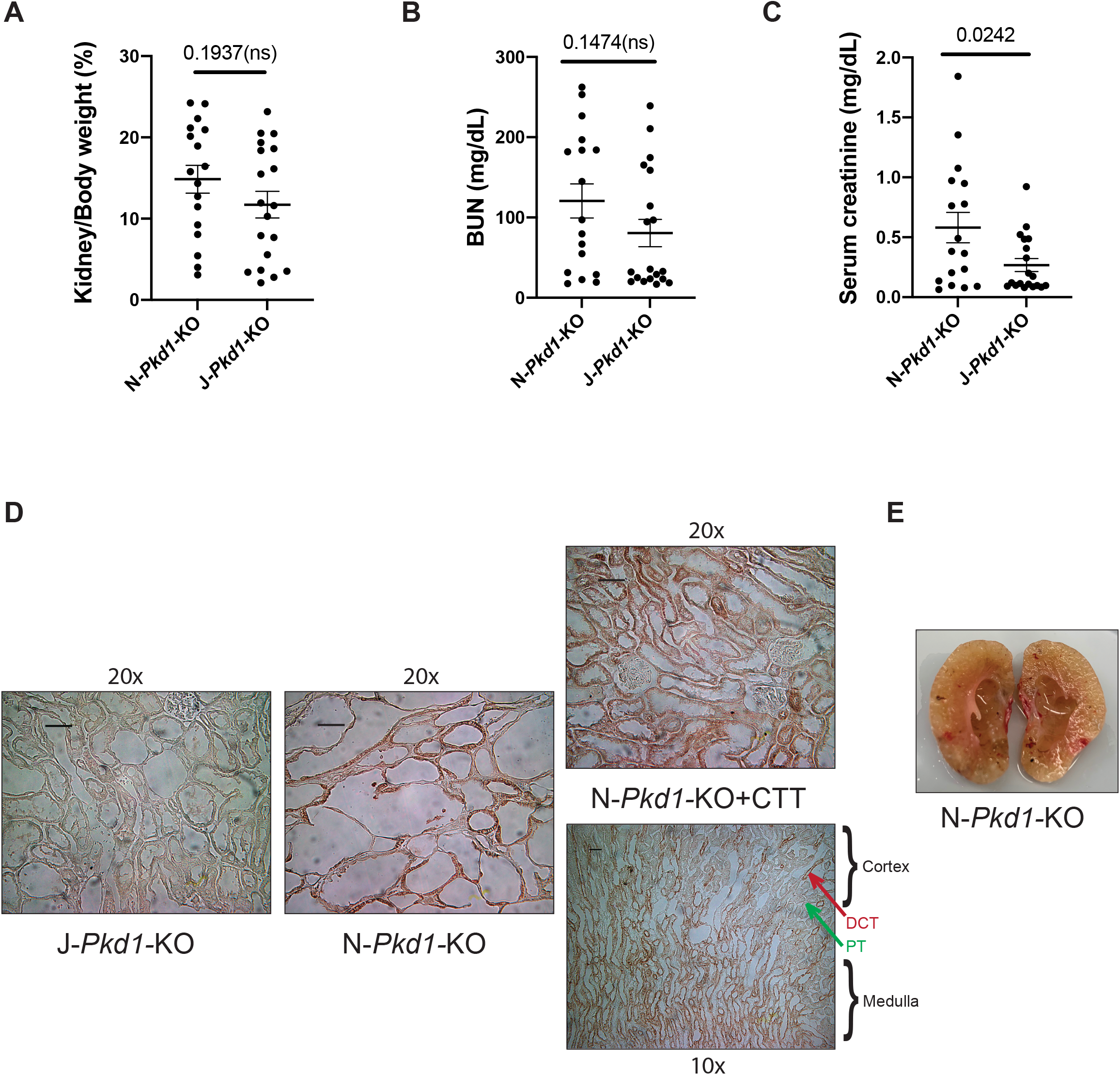
NNT expression alone is not sufficient to suppress the cystic phenotype in *Pkd1*-KO mice. (A-C) Comparative assessment of renal phenotype between N-*Pkd1*-KO and J-*Pkd1-*KO mice, including KW/BW ratio (A), BUN (B) and serum creatinine (C). Data taken from results previously plotted in Figures 1C, D, E and 3A, B, C. Both cystic mouse cohorts are composed of 53% female and 47% male mice. (D) IHC (20X) labeling for NNT in kidney sections of J-*Pkd1-*KO, N*-Pkd1*-KO and N*-Pkd1*-KO+CTT mice acquired by transmission light microscopy and processed with ImageJ automatic settings. IHC image at 10X of N*-Pkd1*-KO+CTT demonstrates the preferential localization of NNT to distal convoluted tubules (DCT) and medullary tubules as compared to proximal tubule (PT). (E) Gross anatomy of a kidney from a 16-week-old N-*Pkd1*-KO mouse, sectioned in its sagittal plane, revealing the presence of moderate hydronephrosis. Scale bar: 50 μm. Data are expressed as mean ± SEM. Pairwise comparisons were performed using Student’s t-test.

### PC1-CTT expression rescues NNT activity

We assessed whether and how the CTT might alter NNT activity. We initially adopted a targeted LC-MS method to directly quantify NAD(P)(H) levels in whole kidney homogenates from 16-week *Pkd1*-KO+CTT and *Pkd1*-KO mice on both the “N” and “J” backgrounds. N-*Pkd1*-KO+CTT exhibited an increase in NADPH/NADP and NADH/NAD^+^ ratios when compared to N-*Pkd1*-KO only mice (Figure 7B and 7C), suggesting CTT-induced effects on the modulation of these cofactors. As expected, expression of CTT in the J-*Pkd1*-KO mice did not affect either ratio. While NAD(P)(H) measurements provide indirect insights into the level of NNT function, these values alone cannot be employed to directly estimate NNT enzymatic activity since a large number of processes contribute to determining NAD(P)(H) levels. Previous studies in non-renal tissue have suggested that 50-70% of the mitochondrial NADPH pool is maintained by isocitrate dehydrogenase (IDH2)(Rydstrom, 2006, Nickel et al., 2015). In light of this potential complexity, we opted to directly evaluate the NNT activity in mitochondrial fractions from N-*Pkd1*-KO+CTT and N-*Pkd1*-KO kidneys. To ensure that the assessment of NNT enzymatic activity was not influenced by the cystic phenotype we conducted this experiment in pre-cystic, 10-week-old mice (Ma et al., 2013)(Figure 7D).

**Figure 7:**
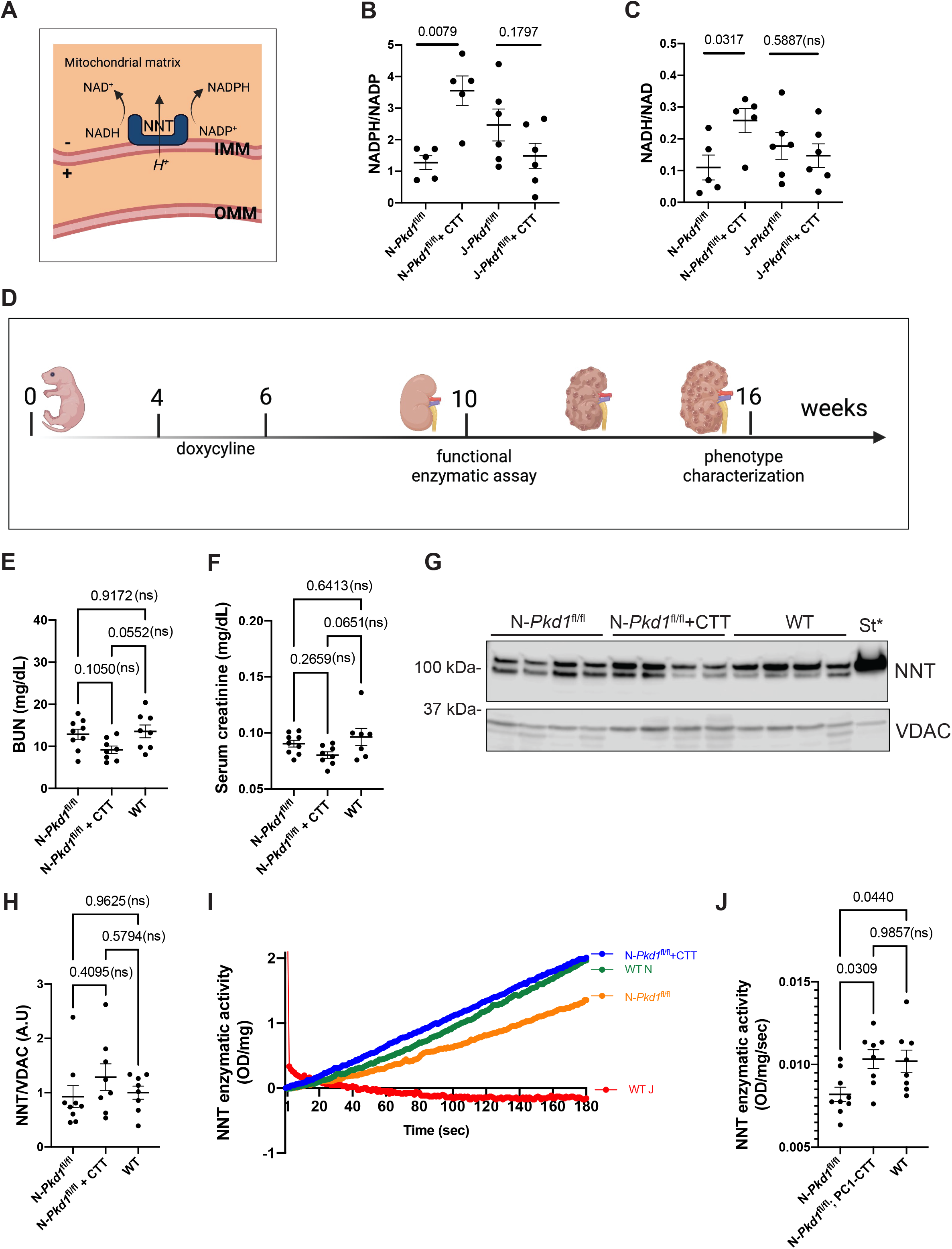
CTT expression modulates mitochondrial redox and increases NNT enzymatic activity. (A) Schematic representation depicting NNT and its localization to the inner mitochondrial membrane and mitochondrial matrix, as well as its dependence on the mitochondrial proton gradient to establish forward enzymatic activity. (B-C) LC-MS detection of NAD(P)(H) cofactors showing significant differences in NADPH/NADP (B) and NADH/NAD^+^ (C) ratios between *Pkd1*-KO+CTT vs *Pkd1*-KO mice exclusively in the “N” background. (D) Schematic representation of experimental timeline for assessment of NNT enzymatic activity in pre-cystic mice at 10 weeks of age. (E-F) Kidney function is preserved in all three 10-week mouse cohorts (N-*Pkd1*-KO, N-*Pkd1*-KO+CTT and N-WT) as shown by normal values of BUN (E) and serum creatinine (F) across the groups. Cystic mouse cohorts are composed of 45%-50% female and 50%-55% male mice. (G-H) Immunoblot of mitochondrial extract from N-*Pkd1*-KO, N-*Pkd1*-KO+CTT and N-WT mouse kidneys (G). *St: mitochondrial extract from rat heart tissue lysate (catalog #ab110341, Abcam) served as standard positive experimental control. VDAC served as mitochondrial loading control. NNT expression (normalized to VDAC) was not significantly different across all groups of 10-week-old mice (H). (I) NNT activity in N-*Pkd1*-KO, N-*Pkd1*-KO+CTT and N-WT mice, quantified using a kinetic spectrophotometric assay that measures reduction of the NAD analog APAD. The measurements were made over the course of 180 s and samples were normalized to protein content. WT “J” mice served as experimental negative controls. (J) Comparison of NNT activity among N-*Pkd1*-KO, N-*Pkd1*-KO+CTT and N-WT mice, measured as ΔOD variation/s/mg of protein. Data are expressed as mean ± SEM. Pairwise comparisons were performed using Mann-Whitney U test due to non-normally distributed data (B and C). Multiple group comparisons were performed using one-way ANOVA followed by Tukey’s multiple-comparisons test (E, F, H and J).

Immunoblotting of a mitochondrial fraction prepared from 70 mg of fresh kidney tissue revealed no significant differences in NNT expression among N-*Pkd1*-KO+CTT, N-*Pkd1*-KO and N-WT mice at 10 weeks of age (Figures 7G and 7H), in marked contrast to differences observed in 16-week-old animals (Figures 5A and 5B). The absence of a significant cystic phenotype in all mice at 10 weeks was consistent with the lack of differences in BUN and serum creatinine (Figures 7E and 7F). The fresh mitochondrial extract was employed in concert with a standard kinetic spectrophotometric assay of NNT enzymatic activity that detects NNT-mediated reduction of the NAD analog APAD (Shimomura et al, 2009). NNT is embedded in the inner mitochondrial membrane and its catalytic domains face the mitochondrial matrix. Thus, to make hydrophilic substrates accessible to the enzyme for the purposes of the assay, the mitochondrial membrane is permeabilized through the addition of detergents Brij35 and lysolecithin. Moreover, a relatively high assay pH is used to minimize nonspecific contributions of NADH-linked dehydrogenases and reductases (Shimomura et al, 2009). We confirmed the specificity of the assay by comparing the activities detected in material prepared from WT “N” to “J” kidneys. While a consistent linear positive slope was observed in assay tracings of material from “N” mice, as expected assay tracings of material from WT “J” mice exhibited no upward slope (Figure 7I). We find that N-*Pkd1*-KO mice exhibit a 20% decrease in NNT enzymatic activity as compared to that detected in healthy “N” WT controls (Figures 7I and 7J). Furthermore, CTT expression in N-*Pkd1*-KO mice rescues NNT enzymatic activity to the same level observed in the healthy “N” controls (Figures 7I and 7J). Of note, this assay was performed on mitochondria extracted from whole-kidney tissue, which includes multiple cell types in addition to Cre-expressing tubular cells. Hence, the magnitude of the observed difference is likely an underestimation of the true effect that is manifest in Cre-expressing cells.

## DISCUSSION

We report the unexpected finding that expressing the C-terminal 200 aa (CTT) of PC1 in an orthologous murine model of ADPKD is sufficient to suppress the development of the cystic phenotype. CTT expression resulted in dramatic preservation of renal function and morphology, as evidenced by BUN and serum creatinine levels that are comparable to levels detected in healthy control mice. Mass spectrometric analysis performed on immunoprecipitates from a crude mitochondrial fraction revealed that NNT is the most significant PC1 binding partner in this setting. We confirmed that NNT interacts with CTT *in vivo* and showed that the CTT and NNT proteins colocalize in mitochondria when CTT is expressed in HEK293 cells. A previous study that employed a split GFP assay to assess PC1-CTT submitochondrial localization showed that PC1-CTT localizes specifically to the mitochondrial matrix or matrix-facing surface of the mitochondrial inner membrane (Lin et al., 2018), a finding consistent with the predicted topological requirements of the novel interaction that we have identified between CTT and the mitochondrial matrix-facing inner membrane enzyme NNT. We showed that the suppression of the cystic phenotype produced by CTT expression is dependent upon this interaction, since no significant rescue is observed when *Pkd1*-KO+CTT mice are compared to *Pkd1*-KO mice in the NNT-deficient “J” background. Similarly, tubular and cystic index and tubular proliferation parameters were only rescued in CTT-expressing *Pkd1*-KO mice generated in the “N” background. Bulk kidney tissue metabolite profiling analysis revealed differentially clustering metabolites only on the “N” background when comparing *Pkd1*-KO+CTT to *Pkd1*-KO mice. We showed that N-*Pkd1*-KO+CTT mice exhibit significant and concomitant downregulation of multiple ADPKD-associated metabolites that have been identified over the past decade and that have led to the suggestion of several metabolism-related potential therapeutic interventions for ADPKD. We assessed the effects of CTT expression on the stable assembly of electron transport chain (ETC) complexes. Previous studies have shown that increased quantities of stably assembled ETC complexes correlate with increased ETC activity and increased levels of oxidative phosphorylation (Sieber et al., 2016, Sing et al., 2014). We show that CTT expression in N-*Pkd1*-KO mice not only leads to increased mitochondrial mass, but also to increased stable assembly of CIV and ATP-synthase after normalization to mitochondrial content. In concert with the metabolomic analysis, which revealed that expression of CTT on the “N” background leads to an approximately 4-fold reduction in lactate levels, these data support the interpretation that the rescue model exhibits a profound shift back towards oxidative phosphorylation serving as the predominant source of ATP generation. Taken together, our data suggest that re-expression of the PC1 C-terminal tail exerts effects that act upstream of the previously identified ADPKD-relevant metabolic pathways.

Interestingly, the presence or absence of NNT expression alone does not dramatically alter the outcome in *Pkd1*-KO mice. In the absence of CTT expression, both “N” and “J” *Pkd1*-KO mice develop severe cystic disease, and our data strongly suggest that the NNT-competent strain exhibits even more aggressive disease progression. While the morphological and physiological consequences of cystic disease are severe in both 16-week-old “J” and “N” *Pkd1*-KO mice, we detect a roughly 2-fold increase in serum creatinine levels in “N” vs “J” *Pkd1*-KO mice. This is especially interesting since declining GFR constitutes a fairly late consequence of cystic disease progression (Grantham et al., 2011). In line with our findings, a recent study characterized the disease progression in a mouse model homozygous for a *Pkd1* hypomorphic variant on three different strain backgrounds: BalbC/cJ (BC), 129S6/SvEvTac (129) and C57BL/6J (Arroyo et al., 2021). While this study does not identify a specific modifier or mechanistic link, it presents robust evidence suggesting that disease progression in C57BL/6J was less severe than in the BC or 129 mice, which are expected to be NNT-expressing. Furthermore, this same study reveals the presence of mild hydronephrosis in both BC and 129 mice but not in C57BL/6J (Arroyo et al., 2021), which is consistent with the frequent presence of hydronephrosis that we observe in N-*Pkd1*-KO mice.

We hypothesize that the more aggressive phenotype in N-*Pkd1*-KO mice may be attributable to NNT’s key role as an antioxidative enzyme due to its capacity to regenerate NADPH from NADP utilizing NADH as the electron donor, driven by the protonmotive force across the mitochondrial inner membrane. By facilitating the detoxification of the reactive oxygen species (ROS) that have been shown to accumulate in hyperproliferative ADPKD tissue (Lu et al., 2020), NNT may allow cystic tissue to overcome oxidative stress and further proliferate. This hypothesis is supported by the trend towards higher proliferation revealed by ki67 staining in “N” vs “J” *Pkd1*-KO mice. In fact, the possible contributions of NNT to the severity of conditions involving hyperproliferation have been recognized in the context of cancer. NNT knockdown, which leads to decreased proliferation and increased apoptosis, is being explored as a therapeutic approach in models of various neoplasms, including gastric cancer (Li et al., 2018), adrenocortical carcinoma (Chortis et al., 2018) and hepatic adenocarcinoma (Ho et al., 2017).

The unexpected finding that NNT expression alone does not protect against cyst formation but that expression of CTT stimulates NNT activity and suppresses cyst formation in an NNT-dependent manner led us to further investigate the mechanism responsible for the NNT-dependent CTT rescue. Our initial efforts involved evaluating potential changes in redox environment secondary to CTT expression. Significant changes in bulk tissue NADH/NAD^+^ and NADPH/NADP ratios were detected between cystic mice expressing or not CTT exclusively in the “N” but not in the “J” background, suggesting that CTT is capable of altering NNT-dependent oxidoreductase pathways. The relative increase of NADPH in the N-*Pkd1*-KO+CTT rescue model suggests the possibility that CTT expression in N-*Pkd1*-KO mice reduces oxidative stress through enhanced NADPH-dependent reduction of oxidized glutathione (GSSG->GSH) and thioredoxin. The impact of oxidative stress on cyst progression in ADPKD models is complex. A previous study suggested that *Pkd1* mutant cells exhibit higher concentrations of oxidized glutathione, which is consistent with increased levels of detrimental oxidative stress (Lin et al., 2018). A different study suggests that exogenous GSH administration may actually enhance cystogenesis by decreasing the vulnerability of mutant cells to ROS and oxidative stress (Flowers et al., 2018). Of note, a previous study has established a connection between NAD^+^ metabolism and ADPKD by showing that administration of nicotinamide led to inhibition of NAD^+^-dependent sirtuin-1 and partially suppressed cyst development in mice (Zhou et al., 2013). These effects were not confirmed in a subsequent patient trial (El Ters et al., 2020).

We further explored the NNT-dependent CTT rescue mechanism by evaluating NNT expression and enzymatic activity. We observed a significant increase in NNT expression in our CTT rescue model compared to cystic N-*Pkd1*-KO mice at the 16-week time point. This finding correlates with the higher quantities of NNT detected in healthy human kidney tissue relative to kidney tissue from ADPKD patients. Interestingly, differences in NNT levels between “N” mice that do or do not express CTT are not observed shortly after *Pkd1* gene disruption, however, there is already significantly greater enzymatic activity in the N-*Pkd1*-KO+CTT rescue mice compared to N-*Pkd1*-KO mice alone at this pre-cystic stage, prior to the development of gross morphological alterations. It is important to note that, to make APAD and NADPH substrates accessible to NNT, it is necessary to disrupt membrane integrity, which eliminates the NNT-driving protonmotive force. Thus, while NNT enzymatic activity can be measured, the driving forces that determine the directionality of the enzymatic reaction remain to be defined. Consequently, we cannot state with certainty whether the effects of CTT expression on NNT ameliorates or exacerbates oxidative damage in cystic cells. It seems likely, however, that the suppressive effects of CTT on cyst development are associated with its capacity to effect NNT-dependent changes in the redox environment. It is, of course, also possible that the observed rescue is not a direct product of NNT enzymatic activity but rather a consequence of NNT-dependent alterations in the mitochondrial protonmotive force. Further experiments will be required to elucidate fully the mechanism of NNT-dependent CTT action.

In summary, we have identified a novel interaction between the mitochondrial enzyme NNT and the C-terminal tail of PC1. More importantly, we showed that expressing the 200 aa C-terminal tail of PC1 in a *Pkd1*-KO murine model of ADPKD is capable of significantly suppressing the cystic phenotype in an NNT-dependent fashion. Recent data shows that re-expression of full-length PC1 in a *Pkd1*-KO murine model results in rapid reversal of ADPKD (Dong et al., 2021). While this finding has broad theoretical and translational implications, the extremely large size of the sequence that encodes the full length PC1 protein severely limits its use in any conventional gene therapy approach. The observation that the PC1-CTT, which is only 200 residues in length, can suppress cyst formation suggests the very exciting possibility that its delivery via gene or protein therapy approaches could impact the course of the disease. It remains to be determined in future studies whether expression of the CTT fragment can recapitulate the ability of the full-length protein to reverse the course of established cystic disease. Whether or not this proves to be the case, the demonstration that the interaction between PC1-CTT and a component of the inner mitochondrial membrane suppresses cyst development opens new directions in the ongoing efforts to both understand the physiological functions of the polycystin proteins and to apply those insights to the development of new therapies.

## ACKNOWLEDGEMENTS

This work was supported by the NIH grants DK120534 (M.J.C. and S.S.), DK072612 (M.J.C.), DoD grant PR191158 (M.J.C. and S.S.); by the Yale School Medicine and Yale West Campus instrumentation support to H.S.; by the Yale School of Medicine startup funds and NIH grant R00 GM124296 to H.S.; and by a PKD Foundation Research Fellowship to L.O. We thank Dr. Hongyu Zhao (Prof. of Statistics and Data Science, Yale University) for generous guidance in designing the statistical analyses performed on all experiments. We thank SueAnn Mentone for outstanding technical assistance with immunofluorescence experiments on mouse tissue, Lonnette Diggs and the George M. O’Brien Kidney Center at Yale (NIH/NIDDK P30 DK079310) for BUN and serum creatinine measurements, and Jie Liu for assisting with tissue metabolite profiling. We thank Raj Pandya for helping with *in vitro* immunofluorescence experiments. We also thank Dr. Alessandra Boletta, Dr. Gerald Shulman, Dr. Leigh Goedeke, Brandon Hubbard and the entire Caplan laboratory for helpful discussions. We are extremely grateful to Drs. Michael Murphy, Luiz Fernando Onuchic and Ana C. Onuchic-Whitford for their critical reviews and valuable contributions to the manuscript. Schematic figures were created with BioRender.com.

## AUTHOR CONTRIBUTIONS

L.O., V.P., and M.J.C. conceived and designed research. L.O., V.P., G.S., V.R., K.D., and X.S. performed experiments. L.O., V.P., G.S., and M.J.C. interpreted data. N.P.G, S.S. and H.S. contributed to experimental design and interpretation. X.S performed bioinformatic analysis of metabolomic data. L.O. and G.S. prepared the figures. L.O. and M.J.C. wrote the manuscript. All authors reviewed the manuscript and contributed valuable inputs.

## DECLARATION OF INTERESTS

Some of the findings presented in this manuscript are included in a provisional patent application filed by Yale University that includes L.O., V.P. and M.J.C. as authors.

## STAR*METHODS

### RESOURCE AVAILABILITY

#### Lead Contact

Further information and requests for resources and reagents should be directed to and will be fulfilled by the lead contact, Michael J. Caplan.

#### Materials Availability

The 2HA-PC1-CTT; *Pkd1*^fl/fl^; Pax8^rtTA^; TetO-Cre mice on both C57BL/6J and C57BL/6N backgrounds were generated in this study and are available through the Yale University School of Medicine.

#### Data and code availability

The mass spectrometry proteomics data have been deposited to the ProteomeXchange Consortium via the PRIDE (Perez-Riverol et al., 2019) partner repository under the dataset identifier PXDXXXXX.

### EXPERIMENTAL MODEL AND SUBJECT DETAILS

#### Mouse models

All animal experiments were approved and conducted in accordance with Yale Animal Resources Center and Institutional Animal Care and Use Committee (IACUC) regulations (protocol # 2019-20088). We utilized the previously characterized *Pkd1*^fl/fl^; Pax8^rtTA^; TetO-Cre (Ma et al., 2013) and *Pkd1*^F/H^-BAC (Fedeles et al., 2011) mouse models. *Pkd1*^fl/fl^; Pax8^rtTA^; TetO-Cre mice were generated on two distinct backgrounds by breeding in either C57BL/6J (stock no: 000664, Jackson Laboratories) or C57BL/6N (stock no:005304, Jackson Laboratories) strains. Additionally, we generated the 2HA-PC1-CTT; *Pkd1*^fl/fl^; Pax8^rtTA^; TetO-Cre on both C57BL/6J and C57BL/6N backgrounds. Animals were maintained at a 12:12 light:dark cycle, with 30-70% humidity and a 20-26°C temperature. In each experiment, animals were age-matched and sex distribution was similar across groups (specific descriptions in figure legends). Cre-negative littermates served as healthy WT controls.

#### Generation of the 2HA-PC1-CTT; *Pkd1*^fl/fl^; Pax8^rtTA^; TetO-Cre mouse model

A *2HA-PKD1-CTT* BAC construct, which encodes a protein corresponding to a 2XHA tag linked to the N-terminus of the final 600 bp of human PC1, was generated with published BAC recombineering technologies (Wang et al., 2009, Warming et al., 2005, Dong et al., 2021). The cDNA sequence encoding *2HA-PKD1-CTT* was introduced into the *pRosa26-DEST* vector (catalog #21189, Addgene) such that a *lox-Neo(R)-3xSTOP-lox* is followed by the sequence encoding *2HA-PKD1-CTT*. A recombination cassette was constructed by flanking a *rpsL+-kana* selection cassette (catalog #20871, Addgene) with two same homology arms (1000bp each arm) from *pRosa26-DEST*. Mouse Rosa26 BAC DNA was electroporated into DY380 bacteria that have stably integrated a defective *λ* prophage containing the Red recombination genes *exo, bet* and *gam* under a strong *pL* promoter controlled by the temperature sensitive *cI857* repressor (kind gift from Dr. Donald Court, National Cancer Institute). The *rpsL+-kana* cassette was introduced into Rosa26 intron 1 region of the BAC after activation of the Red recombination system at 42°C under positive selection by kanamycin resistance. The *rpsL+-kana* cassette in this intermediate was replaced by introducing *lox-Neo(R)-3xSTOP-lox* and *2xHA-PKD1-CTT* fragment with Rosa26 homology arms under negative selection with streptomycin sensitivity conferred by the *rpsL+* gene after the activation of the Red recombination system at 42°C. The final *2xHA-PKD1-CTT Rosa26* BAC was shown to contain only the intended recombination and no other rearrangement using DNA restriction fingerprinting, direct sequencing and *in vitro* recombination in SW106 bacterial strain carrying an L-arabinose-inducible Cre gene. Linearized modified BAC DNA purified by CHEF electrophoresis was used for pronuclear injection to generate transgenic founder lines. The BAC transgenic lines were produced in (C57BL/6J X SJL/J) F_2_ zygotes. Founders were identified by PCR genotyping, verified by sequencing of PCR products and BAC copy number was determined by genomic quantitative PCR as described previously (Dong et al., 2021, Cai et al., 2014, Fedeles et al., 2011). Two BAC founders with BAC copy numbers 2 or 4 were used in this study. All strains were backcrossed at least four generations with C57BL6 and are therefore expected to be at least 90% C57BL6 congenic. These animals were then crossed with *Pkd1*^fl/fl^; Pax8^rtTA^; TetO-Cre mice to generate 2HA-PC1-CTT; *Pkd1*^fl/fl^; Pax8^rtTA^; TetO-Cre, on both C57BL6/J and C57BL6/N backgrounds.

#### Cell Lines

HEK293 cells were cultured in DMEM supplemented with 10% fetal bovine serum (FBS), 1% penicillin/streptomycin, and 1% l-glutamine at 37°C. These cells were then subjected to transient transfection following the protocol described in the transient transfection section of Methods.

#### Human specimens

The Baltimore Polycystic Kidney Disease Research and Clinical Core Center (P30DK090868) provided human kidney tissue from both ADPKD patients and non-affected controls. Samples were surgically harvested according to the guidelines established by the Institutional Review Board of the University of Maryland and were then de-identified. Sex information was not available. Immunoblotting with NNT and actin antibodies was performed using the protocol described in the Western blot section of Methods.

### METHOD DETAIL

#### Mouse kidney tissue harvest

Mice were euthanized according to Yale IACUC protocols. Tail or toe tissue from previously genotyped mice was acquired for a second time when animals were under anesthesia for genotype confirmation. Retro-orbital blood was also collected from these anesthetized mice. The left kidney was excised, weighed, snap-frozen in liquified N_2_ and stored at −80°C for biochemistry analysis. The right kidney was excised, weighed and fixed in 4% paraformaldehyde. Fixed kidneys were then sectioned in half along their sagittal axes, infiltrated with 30% sucrose overnight and embedded in OCT for further imaging.

#### Serum creatinine and BUN measurement

Retro-orbital blood was collected from anesthetized mice prior to sacrifice and centrifuged in Plasma Separator Tubes with Lithium Heparin (BD) to separate plasma. Serum creatinine and BUN analysis were performed by the George M. O’Brien Kidney Center at Yale University.

### Immunoprecipitation

#### Immunoprecipitation from crude mitochondria fractions prepared from *Pkd1*^F/H^-BAC mice

Kidneys from *Pkd1*^F/H^-BAC and WT controls were isolated and homogenized with a Potter Elvehjem homogenizer, followed by a series of differential centrifugations at 4°C according to the following protocol (Wieckowski et al., 2009): lysates were initially submitted to low speed centrifugation at 740g for 5 min, and the collected supernatant was once again submitted to low-speed centrifugation at 740g for 5 min. The collected supernatant from this step was then submitted to high-speed centrifugation at 9,000g for 10 min. The supernatant from this step was discarded and the pellet resuspended and submitted to high-speed centrifugation at 10,000g for 10 min. This final step was repeated twice, and the pellet containing crude mitochondria (both mitochondria and MAM’s) was resuspended in PBS. These samples were incubated with 3mM DTSSP (catalog# 803200-50MG, Sigma) to covalently cross link interacting proteins at room temperature for 30 min in a rocking shaker and then the crosslinking reaction was quenched with 20mM Tris-HCl pH 7.4. Samples were thereafter submitted to a high-speed 10,000g centrifugation for 10 min and resuspended in 100 μl of PBS + 1% SDS, followed by a 30-min immunoprecipitation with 25μl of anti-HA magnetic beads (catalog # 88837, Thermo Fischer Scientific) and 4 washes with TENT buffer (10mM Tris-HCl, 0.1M NaCl, 1mM EDTA, 5% v/v TritonX100). The final immunoprecipitate was either snap frozen in liquified N_2_ and stored at −80°C for further proteomic analysis or eluted in 40μl of 2x Laemmli sample buffer (catalog#1610747, Bio-Rad) with 300mM DTT at 95°C for 10 min for immunoblotting.

Immunoprecipitation from kidney lysates prepared from 2HA-PC1-CTT; *Pkd1*^fl/fl^; Pax8^rtTA^; TetO-Cre mice Kidneys harvested from 2HA-PC1-CTT; *Pkd1*^fl/fl^; Pax8^rtTA^; TetO-Cre mice in both “N” and “J” backgrounds and from *Pkd1*^fl/fl^; Pax8^rtTA^; TetO-Cre mice in the “N” background were snap frozen and stored at −80°C. Homogenization was performed on ice using a motorized tissue grinder (catalog# 1214136, Fisher Scientific) in Tris lysis buffer (50 mM Tris pH 7.4, 100 mM NaCl, 0.5% NP-40, 0.5% Triton X100, 2 mM EDTA) supplemented with complete mini EDTA-free protease inhibitor cocktail tablets (catalog # 11836170001, Roche) and PhosSTOP phosphatase inhibitor cocktail tablets (catalog# 04906837001, Roche). Homogenates were then sonicated for 30 s (2X15 second bursts at 40% power) and incubated for 45 min on ice to complete protein solubilization. Lysates were centrifuged at 8000 rpm for 15 min. Protein concentrations were measured with the Protein Assay Dye Reagent Concentrate (catalog #5000006, Bio-Rad). Anti-HA magnetic beads (catalog # 88837, Thermo Fischer Scientific) were equilibrated in lysis buffer (50 μL beads per reaction in 500μl of lysis buffer) for 10 min at RT on a rocking shaker, and then incubated with a total of 4 mg of tissue lysate per sample, overnight, at 4°C. Following four 5-min washes with 1ml lysis buffer, the dynabeads were magnetically recovered and precipitated proteins were eluted in 60μl of 2x Laemmli sample buffer (catalog#1610747, Bio-Rad) with 300mM DTT. 1/3 of the eluted proteins (20μl) was loaded per sample per gel for immunoblotting.

#### Proteomic analysis

Proteomic analysis on material immunoprecipitated from the *Pkd1*^F/H^-BAC kidneys was performed by the Mass Spectrometry and Proteomics Resource of the W.M. Keck Foundation Biotechnology Resource Laboratory at Yale according to their standard operating procedures. Immunoprecipitated proteins were subjected to chloroform:methanol:water protein extraction, after which they were reduced, alkylated and trypsin digested for follow up LC MS/MS bottom-up data collection. Samples were analyzed with an Orbitrap Fusion mass spectrometer and Mascot Search Engine software was utilized for protein identifications.

#### Western Blotting

Snap-frozen mouse kidneys were homogenized as described in the 2HA-PC1-CTT; *Pkd1*^fl/fl^; Pax8^rtTA^; TetO-Cre whole kidney lysate immunoprecipitation. Protein concentrations were measured with the Protein Assay Dye Reagent Concentrate (catalog #5000006, Bio-Rad). 20-40 μg of protein from whole kidney lysate or 20μl of IP eluted proteins were separated on 4-20% Mini-PROTEAN TGX Precast Protein Gels (catalog # 4561093, Bio-Rad) and electrophoretically transferred to a nitrocellulose membrane. Loading only surpassed the 20-40μg range in the immunoblot depicted in Figure S1, in which loading of 60μg of whole kidney lysate was necessary to identify PC1-CTT in both 2HA-PC1-CTT; *Pkd1*^fl/fl^; Pax8^rtTA^; TetO-Cre and *Pkd1*^F/H^-BAC mice. For western-blotting of human renal tissue, homogenization was performed using a Polytron mechanical homogenizer in Tris lysis buffer (50mM Tris pH 7.4, 100mM NaCl, 0.5% NP-40, 0.5% Triton X100, 1mM EDTA) for 15 s, at 300rpm, on ice. Homogenates were then sonicated for 1 min, with 3 single continuous 15-s bursts at 40% power separated by a 5-s pause, left on ice for 60 min to complete protein solubilization, and centrifuged for 10 min at 10,000g. Membranes were sequentially incubated with blocking buffer (PBS, 6% (w/v) powdered milk/BSA, 0.1% Tween) followed by overnight incubation with primary antibodies. The primary antibodies used in this study were: anti-NNT (#459170, Invitrogen; #sc-390215, Santa Cruz), anti-NNT-HRP (#sc-390236HRP, Santa Cruz), anti-PC1-C-terminus (#EJH002, Kerafast), anti-HA-Peroxidase (#12013819001, Roche), Anti-HA-680 (#26183-D680, Thermo Fischer Scientific), anti-actin (#A2228, Sigma), anti-TOMM20 (# NBP1-81556, Novus Biologicals), anti-Total OXPHOS Cocktail (# MS604-300, Abcam) and anti-VDAC-HRP (#sc-390996HRP, Santa Cruz). All primary antibodies were used at a 1:1000 dilution, except for conjugated primaries anti-NNT-HRP (1:500), anti-HA-Peroxidase (1:500), anti-HA-680 (1:500) and anti-VDAC-HRP (1:250). Unconjugated primary antibodies were detected using species-specific infrared (IR)-conjugated secondary IgG (1:5,000−1:10,000; Li-Cor). Mitochondrial extract from rat heart tissue lysate (#ab110341, Abcam) was utilized as positive control. Membranes were visualized with either the Odyssey Infrared Imager (Li-Cor Biosciences) or Odyssey Fc (Li-Cor Biosciences) for chemiluminescence detection. Individual bands were quantified using ImageJ software (https://imagej.nih.gov/ij/, NIH).

#### Transient transfection in cultured cells

We used Lipofectamine 2000 (catalog# 11668019, ThermoFischer Scientific) to transiently transfect HEK293 cells following the manufacturer’s protocol. Cells were transfected with the previously described 2HA-PC1-CTT construct (Merrick et al., 2019). Briefly, the sequence encoding the final 200 aa of human PC1 (4102-4302) with an N-terminal 2xHA tag was cloned into the pcDNA3.1 zeo vector. The 2HA-PC1-CTT sequence is identical to that expressed in 2HA-PC1-CTT; *Pkd1*-KO mice.

#### Immunofluorescence staining in cells

HEK293 cells grown on poly-L coated coverslips were fixed with 4% PFA in PBS for 30 min at room temperature followed by a 15-min treatment with permeabilization buffer (PBS, 1mM MgCl_2_, 0.1mM CaCl_2_, 0.1% BSA, 0.3 % Triton X100). Cells were then blocked with goat serum dilution buffer (GSDB; 16% filtered goat serum, 0.3% Triton X-100, 20mM NaPi, pH 7.4, 150 mM NaCl) for 30 min, followed by a one-hour incubation with primary antibodies (1:100) diluted in GSDB. The primary antibodies utilized were anti-PC1-C-terminus (catalog #EJH002, Kerafast) and anti-NNT (catalog #459170, Invitrogen). Following 3 PBS washes, samples were incubated with secondary antibodies (1:200) diluted in GSDB for one hour and then washed again with PBS. Alexa Fluor conjugated antibodies (Alexa-594, 647; catalog #A11032 and #A31573 respectively, Life Technologies Invitrogen) were used as secondary reagents. Finally, coverslips were mounted on slides with VectaShield mounting medium (catalog # H-1000-10, Vector Laboratories) and imaged using a Zeiss LSM780 confocal microscope. Images are the product of 8-fold line averaging and contrast and brightness settings were chosen so that all pixels were in the linear range.

#### Mouse tissue immunohistochemistry

Kidneys were fixed and processed for immunohistochemistry as described in the “mouse kidney tissue harvest” section of methods. Four-μm thick sections were heated in 10-mM citrate buffer for 15 min. Slides were then blocked with 0.5% H_2_0_2_ in methanol for 30 min, followed by three 5-min 0.01M PBS washes and further blocking with skim milk in PBS for 1 h at room temperature. Overnight incubation was performed with anti-NNT antibody (catalog# sc-390215, Santa Cruz) at a 1:50 dilution followed by detection using Vectastain Elite ABC-HRP kit (catalog# PK-6200, Vector Laboratories) according to the manufacturer’s instructions.

#### Proliferation assay

Kidneys were fixed and processed for immunofluorescence as described in the “mouse kidney tissue harvest” section of methods. Antigen retrieval for ki67 was performed on 4-μm thick sections by heating slides in a 10-mM citrate buffer for 20 min. After a 30-min incubation with blocking buffer (PBS,1%BSA,10%goat serum), sections were co-incubated with anti-ki67 (catalog# VP-RM04, Vector Laboratories) and anti-Na,K-ATPase α subunit (catalog# a5, DSHB) primary antibodies at a 1:100 dilution followed by detection with Alexa Fluor-conjugated secondary antibodies (Life Technologies Invitrogen) at a 1:200 dilution and Hoechst nuclear staining (catalog #H3570, Molecular Probes Invitrogen). Confocal images were obtained using a Zeiss LSM780 confocal microscope. Images are the product of 8-fold line averaging and contrast and brightness settings were chosen so that all pixels were in the linear range. Anti-Na,K-ATPase α-subunit was used as a tubular marker. Three images were acquired in the upper, middle and lower third of the kidney by a blinded investigator who also quantified the percentage of ki67 positive nuclei relative to total tubular nuclei in these 9 independent images. A total of at least 2000 tubular nuclei were counted per animal.

#### Morphological analyses

Whole kidney images from hematoxylin and eosin–stained sagittal kidney sections were obtained at a 4x magnification using automated image acquisition by the scan slide module in MetaMorph (Molecular Devices). The whole kidney was defined as the region of interest and the ImageJ default auto threshold function was employed to measure cystic and tubular area relative to total kidney area.

#### NNT enzymatic assay

Levels of NNT enzymatic activity were measured in N-*Pkd1*-KO, N-*Pkd1*-KO+CTT and N-WT mice according to a previously established protocol (Shimomura et al, 2009). Briefly, 10-week old pre-cystic (Ma et al., 2013) mice were euthanized according to Yale IACUC protocols. Left kidneys were extracted and partitioned in half in their coronal axis. One half of the kidney (approximately 70mg) was used for mitochondrial preparation, which was performed using the Qproteome^TM^ Mitochondria Isolation Kit (catolog# 37612, Qiagen). The Qiagen protocol was followed in detail until step 11a but the final wash (step 12a) was not performed in order to follow the suggested NNT-assay protocol (Shimomura et al, 2009). Suspension of the final pallet was carried out with 20μl of mitochondria storage buffer (catolog# 37612, Qiagen), a volume sufficient to allow the NNT assay and protein determination. Protein concentrations were measured with the Protein Assay Dye Reagent Concentrate (catalog #5000006, Bio-Rad), revealing a mean concentration of 7.64 +/- 2.2mg/ml. These values were within the protocols’ 5-10mg/ml predicted concentration. The assay medium was prepared and used within 24 h and was composed of 50mM Tris-HCl (pH8.0), 0.5% Brij 35, 1mg/ml of lysolecithin and 300μM of both NADPH and APAD. The assay was performed with 1 ml of assay buffer and 10μl of the mitochondrial suspension and was read with a Bechman DU-640 UV-Vis spectrophotometer with a time-course setting: one measurement per s at a 375nm wavelength, the chosen wavelength for reduced APAD. WT C57BL6/J and C57BL6/N were used as negative and positive controls, respectively. The former confirmed assay specificity (revealing no relevant activity) while the latter confirmed sustained linear activity for several min, as reported in the original protocol (Shimomura et al, 2009). This experiment was repeated three times. The protocol predicts occasional delays in activity initiation, reflecting the time taken for mitochondria to become permeable to substrates, and therefore an investigator blinded to genotype marked the starting point of linear slope. Reactions were measured for a minimum of 150 s of optimal linear slope. The enzymatic activity was calculated by dividing the variation in optical density (OD; y axis) per variation in time (x axis, in s). Results are presented in activity per mg of protein, as established by the protocol (Shimomura et al, 2009).

#### Genomic DNA isolation and quantitative RT-PCR

The DNeasy Blood & Tissue kit was used to extract genomic DNA from all 2HA-PC1-CTT; *Pkd1*^fl/fl^; Pax8^rtTA^; TetO-Cre and *Pkd1*^fl/fl^; Pax8^rtTA^; TetO-Cre mice included in the 16-week cohort, starting with 20mg of kidney tissue from each animal and following the manufacturer’s instructions. Of note, we performed the optional 2-min treatment with 4μl of RNAse A (100mg/ml) at room temperature to obtain RNA-free genomic DNA in transcriptionally active tissues. Quantitative RT-PCR (qRT-PCR) was performed using iTaq Universal SYBR Green Supermix (catalog# 172-5121, Bio-rad). All samples were loaded in triplicates and reactions and data acquisition were performed using the Agilent real-time PCR system with its associated software. GAPDH levels were measured to normalize gene expression. Primers are listed below:

### Determination of Pax8^rtTA^ and TetO-Cre copy numbers

Pax8^rtTA^ F: 5’-AAGTCATAAACGGCGCTCTG-3’

Pax8^rtTA^ R: 5’-CAGTACAGGGTAGGCTGCTC-3’

TetO-Cre F: 5’-TCCATAGAAGACACTGGGACC-3’

TetO-Cre R: 5’-AGTAAAGTGTACAGGATCGGC-3’

GAPDH F: 5’-TGGTGTGACAGTGACTTGGG-3’

GAPDH R: 5’-GTCCTCAGTGTAGCCCAAGA-3’

Mouse samples were normalized to DNA obtained from a control mouse that expressed a single copy of both Pax8^rtTA^ and TetO-Cre. All animals included in the present cohort presented a 1:1 or 2:1 ratio for both genes when compared to controls, confirming homozygosity or heterozygosity for both Pax8^rtTA^ and TetO-Cre alleles.

### Determination of *Pkd1* rearrangement levels

Both forward and reverse primers are situated within the floxed region (exons 2-4) of the *Pkd1* gene in the *Pkd1*^fl/fl^; Pax8^rtTA^; TetO-Cre. The primer sequences used were: *Pkd1* F: 5’-TCTGTCATCTTGCCCTGTTCC-3’ R: 5’-GTTGCACTCAAATGGGTTCCC-3’. The reverse primer is located in Chr17:24,783,583 (exon 4) and the forward primer is contained in the prior intron at position Chr17:24,783,440. The amplified segment is therefore only present in the presence of intact WT *Pkd1*. GAPDH levels were measured to normalize gene expression. All cystic mice were then normalized to the same 4 healthy controls and, as expected, exhibited lower expression of WT *Pkd1* compared to these WT controls. The ratio established between each cystic animal and WT controls served to define the rearrangement levels shown in Figure S3.

### Metabolomics

#### LC-MS metabolite profiling methods

LC/MS-based analyses were performed on a Q Exactive Plus benchtop orbitrap mass spectrometer equipped with an Ion Max source and a HESI II probe, which was coupled to a Vanquish UHPLC (Thermo Fisher Scientific). Polar metabolite extraction and detection methods were adapted from previous literature with minor modifications to be compatible with NAD(P)(H) measurement (Lu et al., 2018, Shen et al., 2017, Wang et al., 2019) (bioRxiv 2021.09.22.461361). Specifically, 40 mg of snap-frozen tissue samples were ground using a mortar and pestle on dry ice. Metabolites were extracted with 800 μl 4/4/2 acetonitrile/methanol/water with 0.1 M formic acid, vortexed and incubated on dry ice for 3 min, and neutralized with 69.6 μl 15% ammonium bicarbonate. The samples were incubated on dry ice for 20 min, then centrifuged at 21,000g for 20 min at 4 °C, and 150 μl of supernatant were transferred to an LC-MS glass vial for analysis.

Polar metabolites were analyzed on Xbrige BEH Amide XP HILIC Column, 100Å, 2.5 μm, 2.1 mm x100 mm (catalog#186006091, Waters) for chromatographic separation. The column oven temperature was 27°C, the injection volume 10 μl and the autosampler temperature 4°C. Mobile phase A was 5% acetonitrile, 20 mM ammonium acetate/ammonium hydroxide, pH 9, and mobile phase B was 100% acetonitrile. LC gradient conditions at flow rate of 0.220 ml/min were as follows: 0 min: 85% B, 0.5 min: 85% B, 9 min: 35% B, 11 min: 2% B, 13.5 min: 85% B, 20 min: 85% B. The mass data were acquired in the polarity switching mode with full scan in a range of 70-1000 m/z, with the resolution at 70,000, with the AGC target at 1e^6^, the maximum injection time at 80 ms, the sheath gas flow at 50 units, the auxiliary gas flow at 10 units, the sweep gas flow at 2 units, the spray voltage at 2.5 kV, the capillary temperature at 310°C, and the auxiliary gas heater temperature at 370°C. Compound Discoverer (Thermo Fisher Scientific) was used to pick peaks and integrate intensity from raw data. The metabolite lists were filtered with minimal peak area > 1e^7^ and annotated by searching against an in-house chemical standard library with 5-ppm mass accuracy and 0.5 min retention time windows followed by manual curation. The data were normalized to tissue protein content. The PCA plots were generated by ClustVis (https://biit.cs.ut.ee/clustvis/); the volcano plots were generated using R script (https://www.r-project.org/).

NAD(P)(H) levels in the kidney extracts were analyzed on SeQuant ZIC-pHILIC polymeric 5 μm, 150×2.1 mm column (EMD-Millipore, 150460). Mobile phase A: 20 mM ammonium carbonate in water, pH 9.6 (adjusted with ammonium hydroxide), and mobile phase B: acetonitrile. The column was held at 27 °C, injection volume 5 μl, and an autosampler temperature of 4°C. LC conditions at flow rate of 0.15 ml/min as following: 0 min: 80% B, 0.5 min: 80% B, 20.5 min: 20% B, 21.3 min: 20%B, 21.5 min: 80% B till 29 min. The data were analyzed using the Xcalibur software.

### QUANTIFICATION AND STATISTICAL ANALYSES

Data quantification and plotting were performed using GraphPad Prism software (https://www.graphpad.com/scientific-software/prism/), with the exception of metabolomic data, which were analyzed and plotted with ClustVis and R as described in the “Metabolomics” section of methods. Data were expressed as means ± SEM. Student’s t-test or Mann-Whitney U test was used for pairwise comparisons, as indicated in the figure legends. One-way analysis of variance (ANOVA) followed by Tukey’s multiple-comparison test was used for multiple comparisons. *P*<0.05 was considered statistically significant. Sample sizes were determined based on our experience working with the *Pkd1^fl^*^/fl^; Pax8^rtTA^; TetO-Cre, and *Pkd1*^F/H^-BAC mouse models; no prior power analysis was performed. Sex distribution was similar between the compared groups. The exclusion criteria were based on mouse well-being; no mice were excluded from this study.

## SUPPLEMENTAL INFORMATION LEGENDS

**Supplemental Figure 1:** CTT expression levels in N-*Pkd1*-KO+CTT mice are comparable to those detected in *Pkd1*^F/H^-BAC mice (A-B) Immunoblot of 60 μg of total kidney lysate from WT, N-*Pkd1*-KO+CTT and BAC-*Pkd1* mice. Actin served as loading control. The 150-kDa bands exclusive to BAC-*Pkd1* mice represent the PC1-CTF fragment that results from N-terminal cleavage of full-length PC1 at the GPS site (Qian et al., 2002). *Pkd1*^F/H^-BAC mice showed the same 37-kDa C-terminal tail fragment band as the CTT-expressing *Pkd1*-KO mice (A), which is present at similar levels of expression (B).

**Supplemental Figure 2:** Pax8^rtTA^ and TetO-Cre copy number do not correlate with disease severity (A) *Pkd1*-KO+CTT and *Pkd1*-KO on the “N” and “J” backgrounds show random distribution of homozygosity or heterozygosity status for both Pax8^rtTA^ and TetO-Cre alleles, and these parameters do not correlate with phenotype severity across any of the four groups.

**Supplemental Figure 3:** PC1 rearrangement levels secondary to Cre-mediated recombination are similar across all cystic mouse cohorts (A) Schematic representation of the generation of qPCR primers capable of exclusively detecting genomic DNA sequence encoding full-length endogenous PC1 from cells that did not undergo Cre-recombination in *Pkd1*-KO mice that do or do not express the CTT protein transgene. Primer positions were based on the mouse genome assembly GRCm39. (B) Comparative analysis of PC1 rearrangement levels across all mouse cohorts.

**Supplemental Figure 4:** Distribution of *Crb1* alleles in cystic mice on the “N” background. (A) The frequency of the *rd8* mutant allele, characteristic of the C57BL/6N background, was similar in N-*Pkd1*-KO+CTT and N-*Pkd1*-KO mice, as determined by standard genotyping (Mehalow et al., 2003).

**Table S1:** Comparative proteomic analysis of material immunoprecipitated from crude renal mitochondrial fractions with anti-HA antibodies from *Pkd1*^F/H^-BAC and WT mice, generated on mixed backgrounds (related to Figure 2C). The table lists the X values (log2 Fold change BAC/control) and Y values (-log10 *P* value), as depicted in the volcano plot in Figure 2C, for all identified peptides and indicates the targets with *P* value <0.05.

**Table S2:** Untargeted metabolomic analysis comparing lysates of kidneys from *Pkd1*-KO and *Pkd1*-KO+CTT mice on the “N” and “J” backgrounds (related to Figure 4). The table lists metabolites detected by mass spectrometry analysis as well as the comparative analysis with fold change and *P* values performed on both backgrounds.

## Notes

### Competing Interest Statement

The authors have declared no competing interest.

