## Supplementary figures and images for "The C-terminal tail of polycystin-1 suppresses cystic disease in a mitochondrial enzyme-dependent fashion"

### Figure S1

#### Supplemental Figure 1

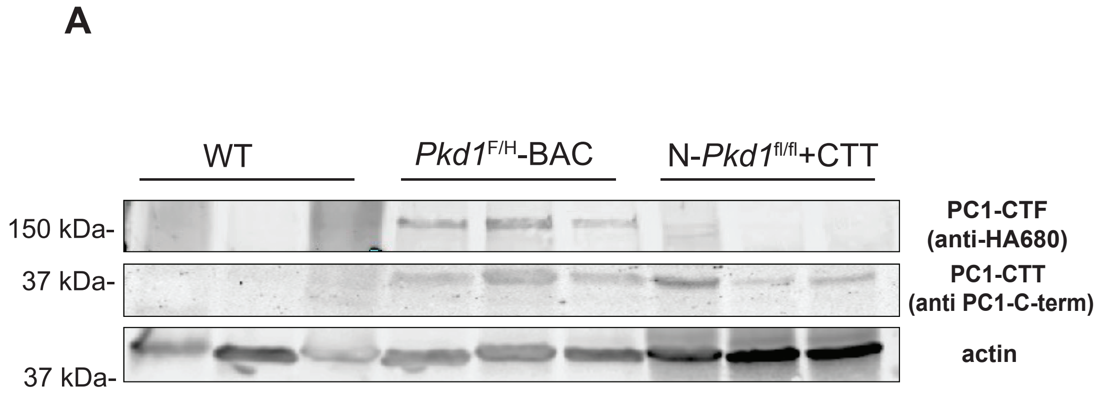


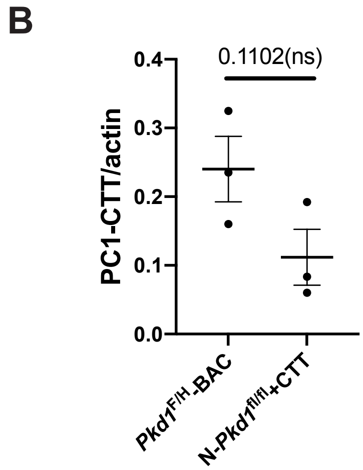

### Figure S2

#### Supplemental Figure 2

**A**

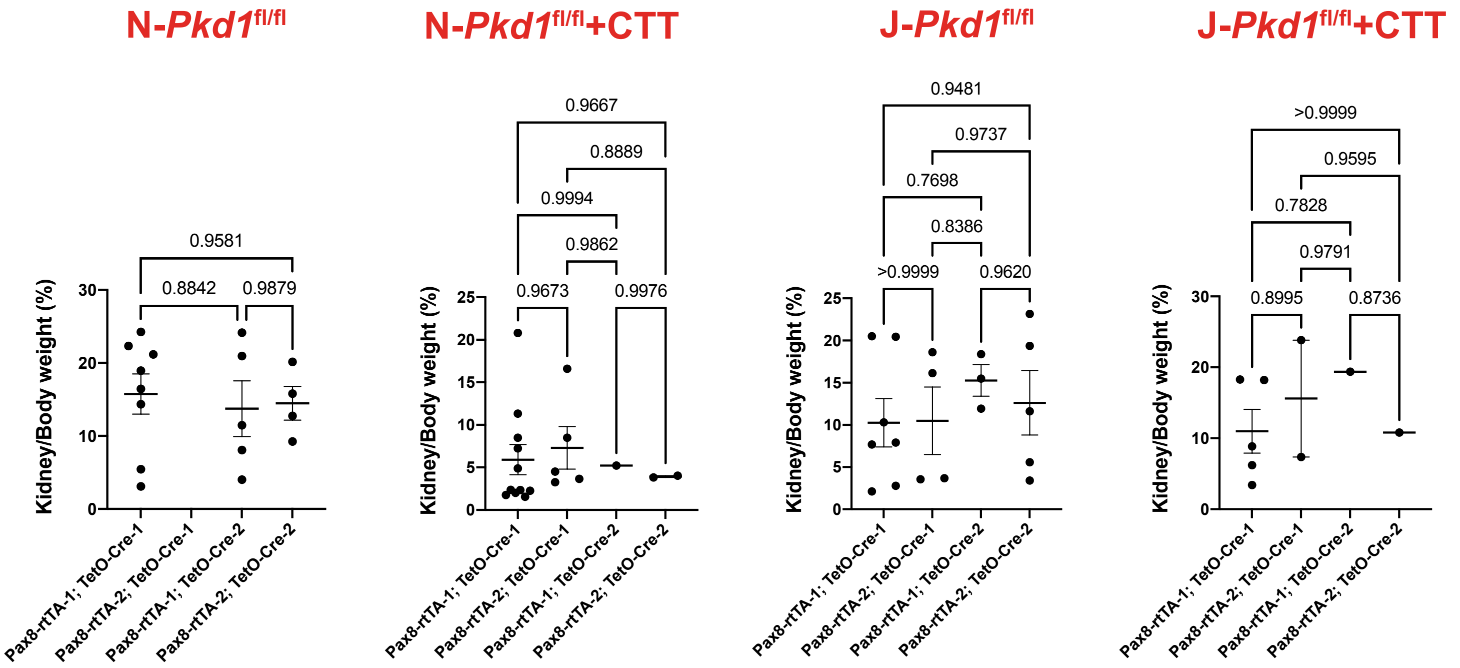

### Figure S3

#### Supplemental Figure 3

A

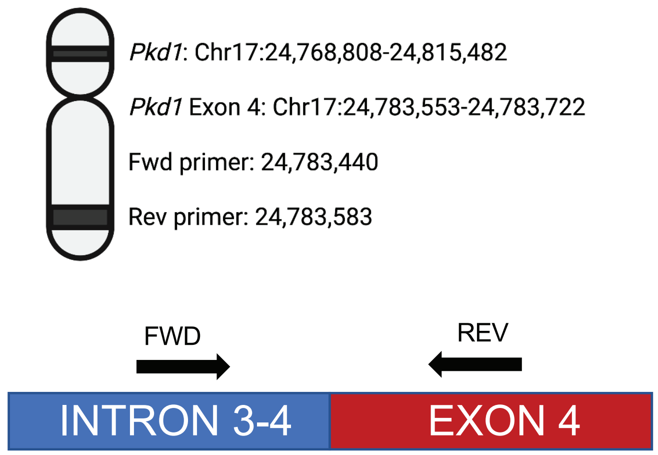


B

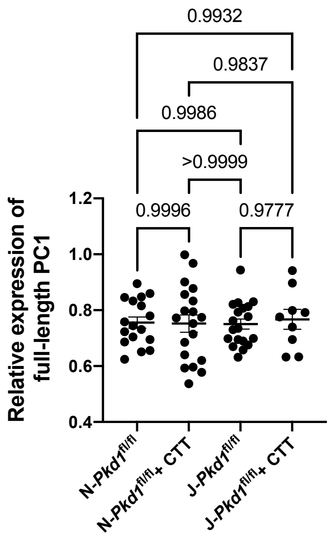

### Figure S4

#### Supplemental Figure 4

A

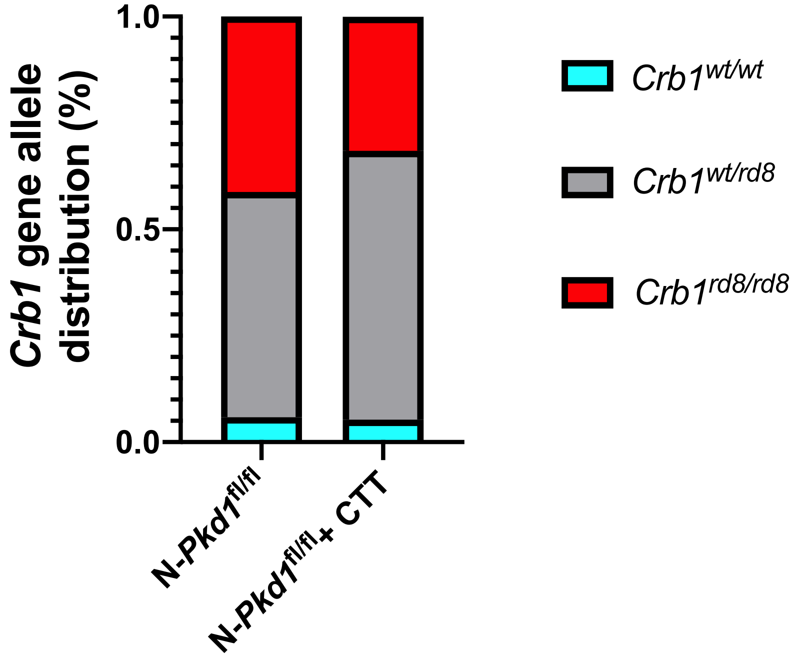
